# Clinical interpretation of integrative molecular profiles to guide precision cancer medicine

**DOI:** 10.1101/2020.09.22.308833

**Authors:** Brendan Reardon, Nathaniel D Moore, Nicholas Moore, Eric Kofman, Saud Aldubayan, Alexander Cheung, Jake Conway, Haitham Elmarakeby, Alma Imamovic, Sophia C. Kamran, Tanya Keenan, Daniel Keliher, David J Konieczkowski, David Liu, Kent Mouw, Jihye Park, Natalie Vokes, Felix Dietlein, Eliezer M Van Allen

**Author notes:** **CORRESPONDING AUTHOR**: Eliezer M Van Allen.

## Abstract

Individual tumor molecular profiling is routinely used to detect single gene-variant (“first-order”) genomic alterations that may inform therapeutic actions -- for instance, a tumor with a *BRAF* p.V600E variant might be considered for RAF/MEK inhibitor therapy. Interactions between such first-order events (e.g., somatic-germline) and global molecular features (e.g. mutational signatures) are increasingly associated with clinical outcomes, but these “second order” alterations are not yet generally accounted for in clinical interpretation algorithms and knowledge bases. Here, we introduce the Molecular Oncology Almanac (MOAlmanac), a clinical interpretation algorithm paired with a novel underlying knowledge base to enable integrative interpretation of genomic and transcriptional cancer data for point-of-care treatment decision-making and translational hypothesis generation. We compared MOAlmanac to first-order interpretation methodology in multiple retrospective patient cohorts and observed that the inclusion of preclinical and inferential evidence as well as second-order molecular features increased the number of nominated clinical hypotheses. MOAlmanac also performed matchmaking between patient molecular profiles and cancer cell lines to further expand individualized clinical actionability. When applied to a prospective precision oncology trial cohort, MOAlmanac nominated a median of two therapies per patient and identified therapeutic strategies administered in 46% of patient profiles. Overall, we present a novel computational method to perform integrative clinical interpretation of individualized molecular profiles. MOAlmanc increases clinical actionability over conventional approaches by considering second-order molecular features and additional evidence sources, and is available as an open-source framework.

## INTRODUCTION

Targeted panels or whole-exome sequencing now routinely inform the clinical care of oncology patients^1^. The resulting collections of patient-specific cancer genome alterations are valuable resources in the advancement of precision medicine. However, the growing quantity and complexity of potentially actionable genomic alterations available for each patient limit the ability of any individual clinician or researcher to interpret them. This challenge necessitated the creation of clinical interpretation algorithms to computationally prioritize large sets of patient-specific alterations by clinical and biological relevance, as well as exposed the need to pair these interpretation algorithms with up-to-date knowledge bases that link molecular alterations to relevant clinical actions.

Clinical decision-making in precision oncology commonly emphasize “first-order” relationships -- pairing individual somatic variants, copy number alterations, pathogenic germline variants, or fusions with specific clinical actions such as use of *BRAF* p.V600E and RAF/MEK inhibition -- based on FDA approvals and other clinical evidence^2–7^. While these efforts have been highly fruitful, they also have certain limitations. Many academic and commercially available targeted panels focus primarily on somatic variants and copy number alterations; often, they do not sequence associated germline tissue or comprehensively assess fusions^1^. Yet pathogenic germline variants impact cancer risk and can also modify clinical interpretation of secondary somatic events in the same gene or of genome-wide mutational signatures, e.g. DNA repair^8,9^. Similarity, the approval of TRK inhibitors for patients with any solid tumor harboring *NTRK* fusions and other biological insights gained from somatic variants that can be identified from RNA may warrant expanding routine clinical sequencing to jointly evaluate a patient’s genomic and transcriptional data^10,11^. In addition, the ongoing characterization of the cancer genome has revealed the importance of considering these first-order events in tandem as well as “second-order” molecular features -- genomic processes such as microsatellite instability and tumor mutational burden that are global rather than limited to individual gene(s). Such processes have also been associated with clinical phenotypes, such as COSMIC Signature 6 correlating with mismatch repair deficiency (MMR) and microsatellite instability (MSI) linked to cancer immunotherapy response^12^. Lastly, even with the consideration of these additional features and second-order relationships, some patients may be variant-negative and thus may not qualify for genomically guided treatment. To address this challenge, multiple efforts have demonstrated that preclinical cell line models can also inform treatment selection, but such approaches are constrained by both the limited molecular diversity of cancer cell lines and computational difficulty in matchmaking, to identify which models are most representative of an individual patient’s tumor^13–17^.

To maximize interpretability of integrative molecular profiling for point-of-care treatment decision-making and translational hypothesis generation, new methodologies are needed to leverage both first- and second-order molecular alterations, relationships between multiple co-occurring events, and the full spectrum of both clinical and preclinical evidence. Here, we introduce Molecular Oncology Almanac (MOAlmanac), a clinical interpretation algorithm paired with an alteration-action database (Figure 1) that operates on germline, somatic, and transcriptional data in tandem from individual patients. MOAlmanac expands the scope of considered molecular alterations beyond somatic variants and copy number alterations to include fusions, germline variants, and concordance between events across feature types. In addition, MOAlmanac considers global “second-order” molecular features and introduces a patient-to-cell line matchmaking module to leverage cell line profiling to nominate additional genomic features potentially associated with therapeutic sensitivity. MOAlmanac is provided in a cloud-based framework and delivers reports at the level of the individual patient. By integrating diverse data sources with higher-order interpretation, MOAlmanac expands the landscape of clinical actionability to facilitate point-of-care decision making and to advance precision cancer medicine.

**Figure 1.**
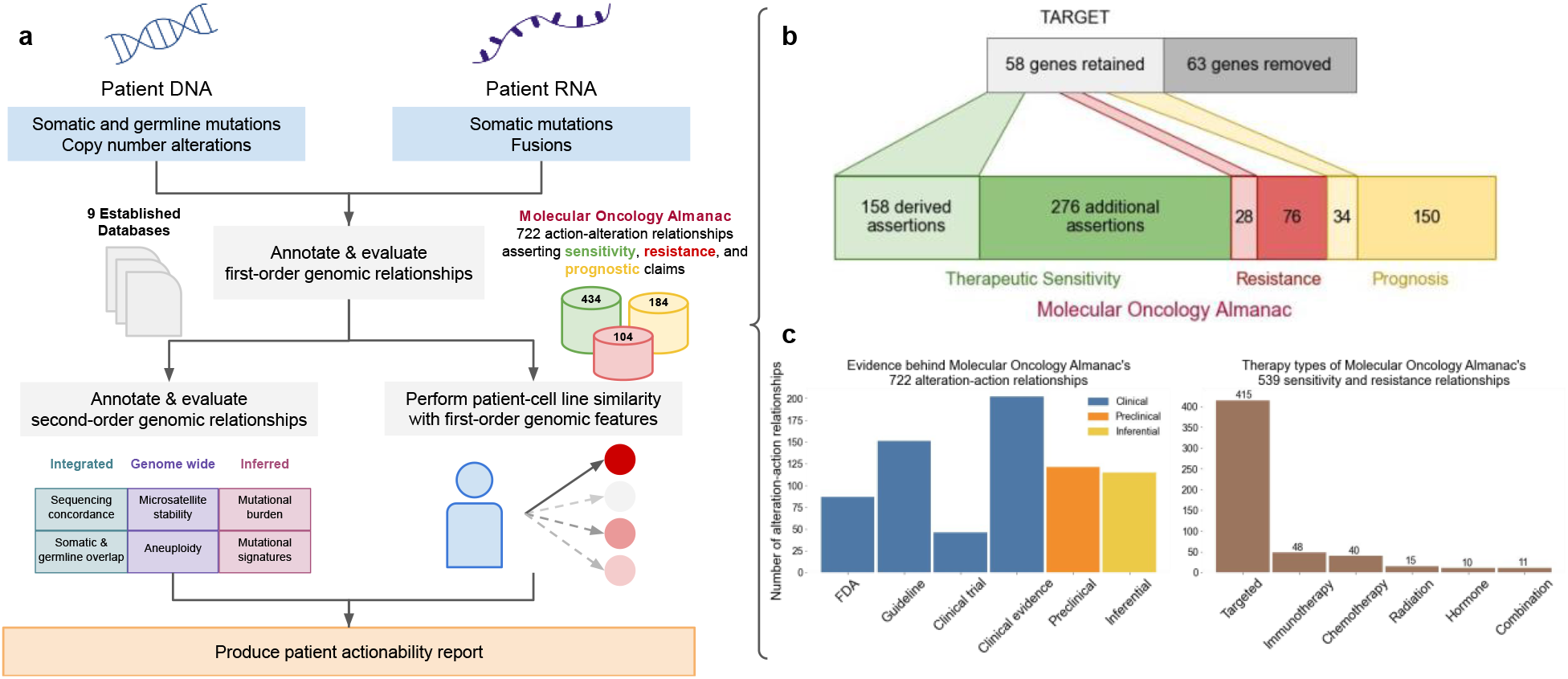
Molecular Oncology Almanac, a clinical interpretation framework. **(a)** The Molecular Oncology Almanac accepts any combination of somatic single nucleotide variants (snvs), insertions and deletions (indels), copy number alterations (cnas), germline snvs and indels, somatic snvs from orthogonal sequencing, and rearrangements from RNA. Molecular features are annotated for clinical relevance and with several other data sources before being heuristically sorted (first-order). Variants are used to evaluate genomic features; somatic-germline overlap, concordance of somatic variants with orthogonal sequencing, COSMIC mutational signature contributions, mutational burden, and MSI related variants (second-order). Somatic mutations, copy number alterations, and fusions are used to assess similarity to individual cell lines for further therapeutic sensitivity suggestions. A report of putative actionability is generated (Methods). **(b)** A literature review was performed to identify relationships between molecular alterations and clinical actions for precision oncology, beginning with relationships suggested in TARGET^2^. 63 genes were removed from TARGET due to insufficient evidence and 58 were retained. Clinical relationships were cataloged as suggesting therapeutic sensitivity, resistance, or prognostic value in an SQL database (Methods) and made available online (https://moalmanac.org). (**c**) Sources catalogued in the Molecular Oncology Almanac, categorized by evidence (left) and therapy types (right)

## RESULTS

### Developing an integrated interpretation framework

Molecular Oncology Almanac is a clinical interpretation method that evaluates individual patient molecular profiles to facilitate precision oncology (Figure 1a). Individual genomic events are annotated and sorted to identify those that are both highly associated with cancer and associated with treatment response or prognosis. First, features are prioritized based on an association between the involved genes and cancer in several data sources; in order: MOAlmanac’s database (described below), Cancer Hotspots, 3D Cancer Hotspots, Cancer Gene Census (CGC), Molecular Signatures Database, and Catalogue of Somatic Mutations in Cancer (COSMIC) (Methods, Supplementary Figure 1a)^18–23^. Next, molecular features are further prioritized based on associations between specific alterations and each data source. For instance, *KRAS* p.G12A ranks higher than *KRAS* p.I36M as both protein changes are reported as 3D hotspots but only p.G12A matches to Cancer Hotspots.

The clinical relevance of each cancer-associated molecular feature is further assessed based on an underlying custom knowledge base, which contains 722 assertions relating molecular features to therapeutic sensitivity, resistance, and prognosis based on published literature and guidelines. This resource evolved from our prior actionability database (Tumor Alterations Relevant for GEnomics-driven Therapy (TARGET)), which represented entries as genes and data types^2^ (Figure 1b, Methods). In contrast, MOAlmanac defines molecular features broadly to encompass the varying types of alterations backed by cited evidence. For example, MOAlmanac is capable of recording information regarding specific singleton features (e.g. *BRAF* p.V600E) but also more general event classes (such as the presence of an *ALK* fusion without regard to the fusion partner). Relationships between molecular features and treatment response are annotated for targeted therapies (415 assertions), immunotherapies (48), chemotherapies (40), radiation therapy (15), hormonal treatments (7), and combination therapies (11) (Figure 1c, Methods). Individual genomic events that match catalogued features are labeled by the specificity of the underlying event and match completeness. For example, exact matches to fully defined features, such as *BCR-ABL1*, are labeled as “Putatively Actionable”; partial matches within a feature type are labeled as “Investigate Actionability”, such as an *ATM* missense variant matching to a catalogued *ATM* nonsense variant; and events whose gene appears in the database under a different data type are highlighted as “Biologically Relevant” but not associated with a clinical assertion, e.g. a *CDKN2A* somatic variant matching to *CDKN2A* copy number deletions. These assertions are derived from numerous evidence sources in accordance with existing frameworks^3–5,24^, including: FDA approvals (FDA-approved), clinical guidelines (Guideline), results from prospective clinical trials (Clinical trial), results from human studies other than a clinical trial (Clinical evidence), findings from cancer cell lines or animal models (Preclinical), or inferences from mathematical models or associations between molecular features (Inferential) (Figure 1c, Methods).

MOAlmanac also characterizes individual features in concert with each other and second-order genomic events. For each MOAlmanac gene, events across all feature types are reported together to elucidate contributions from distinct types of genomic events. Somatic variants in a given gene will increase in priority if either a truncating or a pathogenic or likely pathogenic (according to ClinVar) germline variant appears in the same gene or if the somatic variant is observed with sufficient power in validation sequencing, if provided^24,25^. Both COSMIC mutational signature contributions and tumor mutational burden (TMB) are calculated and variants related to microsatellite instability are highlighted. Tumor ontology is mapped with Oncotree. Tumor purity, ploidy, whole-genome doubling, and microsatellite stability status are also accepted for reporting and evaluation. All nominated clinical associations are reported in a web-based actionability report (Methods).

### Evaluating expanded molecular profiling and actionability in two retrospective cohorts

We first evaluated MOAlmanac relative to our prior established whole-exome sequencing (WES) first-order interpretation framework (PHIAL with TARGET), which considers somatic variants and copy number alterations^2^. WES and RNA-sequencing (RNA-seq) data were acquired for 110 previously published metastatic melanomas (n = 44 with RNA)^26^ and 150 patients with metastatic castration-resistant prostate cancers (mCRPC, n = 149 with RNA)^27^. All samples were analyzed to call somatic variants, germline variants, and copy number alterations from WES and somatic variants and fusions from RNA-seq (Methods).

We compared how often the two methods observed a clinically relevant event associated with therapeutic sensitivity, resistance, or prognosis when only somatic variants and copy number alterations were considered. Furthermore, we characterized only well-established relationships by restricting our analysis to assertions curated from FDA approvals, clinical guidelines, clinical trials, or clinical evidence. MOAlmanac identified 312 such putatively actionable events from 191 patients (73 melanoma, 118 mCRPC), 218 (69.87%) of which were flagged by PHIAL for clinical relevance. For example, the most commonly flagged features were *BRAF* p.V600E (39 patients), *MET* amplification (9), and *PTEN* deletion (9) for metastatic melanomas and *AR* amplifications (82), *PTEN* deletions (40), and *RB1* deletions (21) in mCRPC. When “Investigate Actionability” variants were included, an additional 54 patients (20.8% of cohort) harbored a potentially clinically relevant variant, such as *NRAS* p.Q61K (10, melanoma) with associated sensitivity to selumetinib, 31 of which were also highlighted by PHIAL. PHIAL identified 0 events as Putatively Actionable and 113 as Investigate Actionability which were not highlighted by MOAlmanac; however, all genes associated with these events were not migrated to MOAlmanac from TARGET for reasons such as insufficient evidence of clinical relevance (Methods).

Next, while still limiting our analysis to somatic variants and copy number alterations, we investigated how the inclusion of preclinical and inferential evidence sources affected identification of potentially actionable results. On the basis of preclinical evidence, 120 such genomic events from 107 patients were identified -- for example, *PTEN* deletions and sensitivity to everolimus or AZD8186, 86 (71.7%) of which were also highlighted by PHIAL. Inferential evidence highlighted 19 additional putatively actionable copy number alterations from 19 patients, most prominently *CCND1* amplifications for reported sensitivity to palbociclib (n=15). Thus, using all catalogued evidence, MOAlmanac noted 1175 somatic variants and copy number alterations as Putatively Actionable or Investigate Actionability across 249 patients (109 melanoma, 140 CRPC). Of these events, PHIAL highlighted 73 (6.2%) as Putatively Actionable, 352 (30%) as Investigate Actionability, and 369 (31.4%) as Biologically Relevant.

We then evaluated whether an expanded set of molecular features (including germline variants and fusions as additional first-order features and tumor mutational burden, mutational signatures, and aneuploidy as second-order features, none of which are handled by PHIAL, could further broaden the actionability landscape for individual patients (Figure 2b). Pathogenic and likely pathogenic germline variants highlighted 10 additional clinically relevant molecular features across 10 different samples (0 melanoma, 10 mCRPC), six of which were *BRCA1/2* variants. MOAlmanc identified 127 clinically relevant fusions across 82 patients; ten mCRPC tumors harbored no putatively actionable somatic variants or copy number alterations but did contain *TMPRS22-ERG*. Regarding second-order molecular features, elevated TMB was noted for 43 patients with metastatic melanoma and 4 with mCRPC (Methods), clinically relevant mutational signatures were observed in 40 molecular profiles, and whole-genome doubling, which has been associated with poor prognosis, was observed in 137 profiles^28^. In some of these cases, combinations of these features were particularly relevant when present in tandem. For example, a pathogenic *BRCA2* variant, p.S1882*, was observed in one patient along with a 39% mutational signature attribution to COSMIC Signature 3, both of which may suggest homologous recombination repair deficiency (HRD) and sensitivity to PARP inhibition^29–31^. By considering these feature types, MOAlmanac identified an additional 397 clinically relevant molecular features in 214 patients, resulting in 258 patients with at least one event associated with therapeutic sensitivity, resistance, or prognosis.

**Figure 2.**
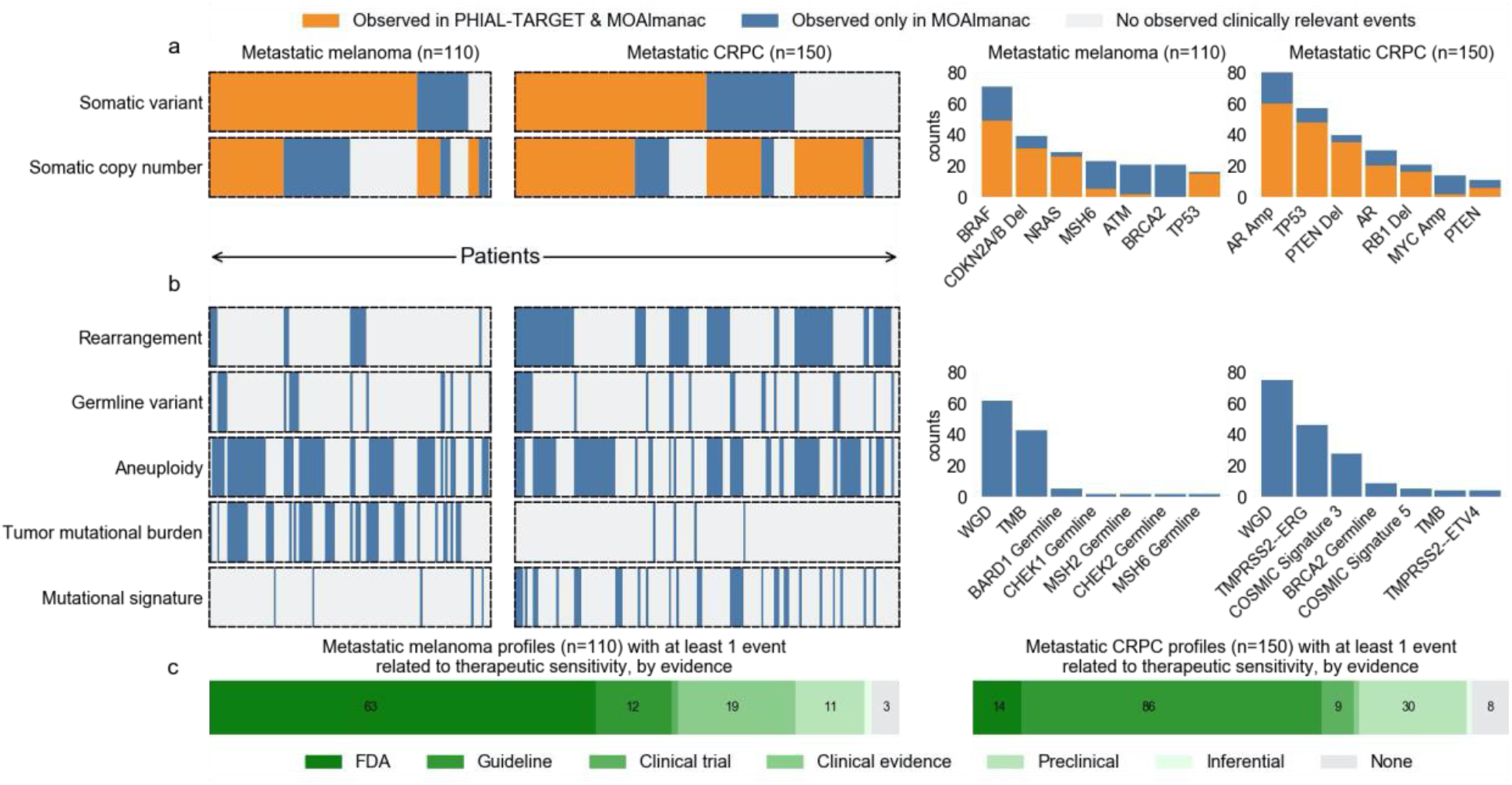
Benchmarking MOAlmanac against PHIAL & TARGET. Molecular Oncology Almanac was benchmarked against PHIAL & TARGET using 110 metastatic melanomas and 150 metastatic castration-resistant prostate cancers^2,26,27^. **(a)** The Molecular Oncology Almanac increased the number of patients with a somatic variant or copy number alteration labeled as “putatively actionable” or “investigate actionability” from 115 to 249 relative to PHIAL; patients are aligned across feature types vertically (left). Specific molecular features that were observed by both PHIAL & MOAlmanac (orange) and by MOAlmanac only (blue) for each cohort are shown (right). **(b)** Features not routinely used in clinical sequencing were utilized to characterize actionability: rearrangements, germline variants, aneuploidy, mutational burden, and mutational signatures; patients aligned with (A) vertically (left). Considering these features types further identified 7 patients with a clinically relevant feature. Specific molecular features that were additionally observed in each cohort are shown (right). (Abbreviations used: WGD = whole genome doubling, TMB = tumor mutational burden). (**c**) Including preclinical evidence when considering putative actionability provides an additional 41 patients (11 patients with metastatic melanoma and 30 patients with castration resistant prostate cancers) with a molecularly matched therapeutic hypothesis.

Focusing specifically on therapeutic sensitivity, such consideration of an extended set of feature types and additional evidence sources provided otherwise variant-negative patients with clinical hypotheses (Figure 2c, Supplementary Table 1). FDA approved or clinical guideline associations resulted in a highlighted therapy for 175 of 260 patients (75 and 100 for melanoma and CRPC, respectively); 11 patients obtained a therapeutic hypothesis from feature types other than somatic variants and copy number alterations, such as elevated TMB (2 patients) or *NTRK* fusions (1). Inclusion of preclinical and inferential evidence sources further decreased the number of variant-negative patients from 85 to 11 (41 preclinical, inferential); for example *CDKN2A/B* deletions and sensitivity to EPZ015666 (6).

In total, MOAlmanac found at least one clinically relevant feature in 100% and 98.6% of metastatic melanoma and mCRPC profiles, using evidence ranging from FDA approvals to inferential relationships and both first- and second-order molecular features (Figure 2a, 2b). In comparison, PHIAL identified such somatic variants and copy number alterations in 92.7% and 89.3% of metastatic melanoma and mCRPC profiles, respectively. Thus, the inclusion of additional feature types and evidence for clinical interpretation provided patients with an expanded set of clinical hypotheses.

### Leveraging preclinical models for clinical actionability

We next investigated whether preclinical data from high-throughput therapeutic screens of cancer cell lines could further inform clinical interpretation within the MOAlmanc methodology. We identified 452 solid tumor cell lines from the Cancer Cell Line Encyclopedia (CCLE) and Sanger Institute’s Genomics of Drug Sensitivity in Cancer (GDSC) that had available data on nucleotide variants, copy number alterations, fusions, and drug sensitivity (Methods)^32,33^. Of MOAlmanac’s 124 catalogued therapies, 44 were represented in the current GDSC2 dataset and 15 additional therapies were represented only in the older GDSC1 dataset. These 44 therapies are involved in 159 catalogued assertions between genomic alterations and therapeutic sensitivity, for each MOAlmanac evaluates sensitivity for wild-type cell lines vs those harboring the corresponding or related alterations. For example, in the case of the catalogued preclinical relationship between *PIK3CA* p.H1047R and sensitivity to pictilisib, MOAlmanac reports sensitivity for wild-type cell lines versus those harboring any genomic alteration in *PIK3CA*, any nonsynonymous variant in *PIK3CA*, any missense variant in the gene, and those specifically with the p.H1047R variant (Supplementary Figure 3a). Across all evaluable relationships asserting sensitivity, 12 therapies showed a significant difference in IC50 between wild type and mutant cell lines (Supplementary Table 2, Methods). Thus, high-throughput therapeutic screens of cancer cell lines are used as an orthogonal axis of evidence to evaluate clinically relevant relationships nominated by MOAlmanac.

The above approach simplistically compares sensitivity between cell lines that do or do not share a single specific molecular feature. A potential limitation of this approach is that it includes cell lines that share the index feature but are otherwise genomically highly dissimilar and therefore whose overall biological relevance to the underlying patient sample may be questionable. Therefore, we were motivated to identify cancer cell lines that shared more extensive similarities in their molecular profiles and investigate whether such “patient-to-cell line matchmaking” could identify additional potential therapeutic sensitivities. Previous approaches have evaluated genomic similarity based on shared mutated genes that are weighted by their recurrence in TCGA^15,16^; however, we chose to assess models based on shared therapeutic sensitivity independent of histology-specific priors. We evaluated several models on cell lines using a hold-one-out approach (Methods). For each cell line, we determined whether its nearest neighbor shared drug sensitivity to any GDSC therapy (Figure 3a, Methods). Similarity Network Fusion applied to nucleotide variants, copy number alterations, and rearrangements involving CGC genes and genomic alterations associated with FDA approvals most frequently assigned a nearest neighbor that shared drug sensitivity (19.7%, Figure 3b, Methods)^34^.

**Figure 3.**
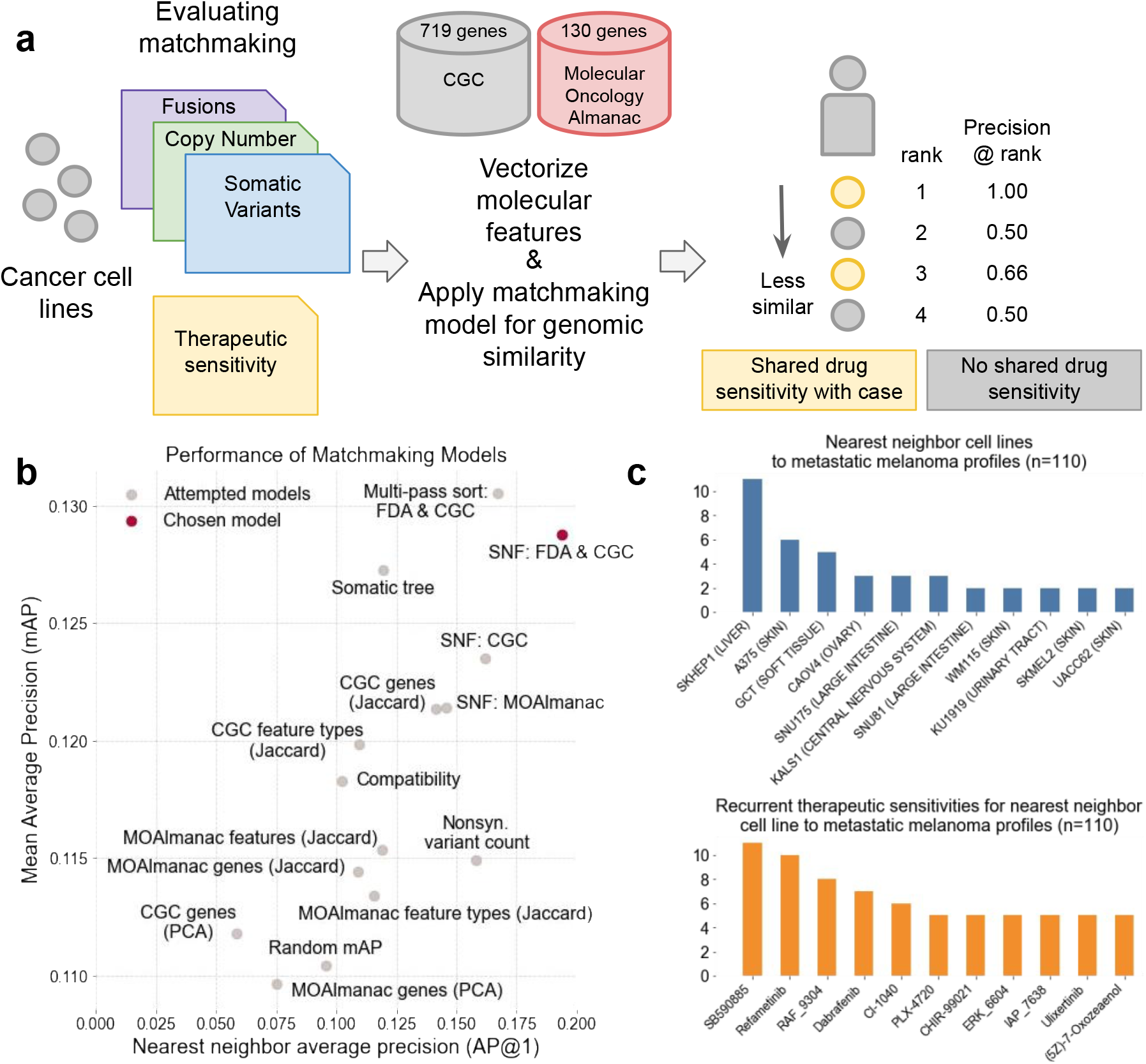
Leveraging preclinical models in MOAlmanac. MOAlmanac leverages preclinical data from cancer cell lines which have been molecularly characterized and subject to high-throughput therapeutic screens to provide supplemental hypotheses through profile-cell line matchmaking. (**a**) Somatic SNVs, CNAs, and fusions of cancer cell lines are formatted, annotated with MOAlmanac and CGC, and vectorized into sample x feature boolean dataframes. Feature sets and similarity metrics were evaluated by their ability to sort cell lines relative to one another based on shared genomic features, such that cell lines that shared therapeutic sensitivity were deemed more similar. Metrics from information retrieval were used for evaluation; mean average precision (mAP, how the model does overall at sorting cell lines which share therapeutic sensitivity to be closer to the case profile) and average precision at rank 1 (ap@1, how often the nearest neighbor shared therapeutic sensitivity). (**b**) Models were evaluated on 377 cancer cell lines using a hold-one-out approach. The model which had the strongest trade off between the two metrics used Similarity Network Fusion to fuse networks of somatic variants, copy number alterations, and fusions in CGC genes with specific MOAlmanac features associated with an FDA approval^21,34^. (**c**) Recurrent nearest neighbors and their sensitive therapies for 110 metastatic melanomas. SKHEP1_LIVER was the first neighbor for 11 profiles, A375_SKIN for six, and GCT_SOFT_TISSUE for five. Nearest neighbors were sensitive to MEK and RAF inhibitors: SB590885 (BRAF inhibitor, 11 neighbors), Refametinib (MEK, 10), RAF_9304 (RAF, 8), and Dabrafenib (BRAF, 7).

This patient-to-cell line matchmaking module was then applied to our previously characterized cohorts of patients with mCRPC and metastatic melanoma. Within the mCRPC cohort, the most common nearest neighbor cell line among the 452 tested was VCaP, one of two prostate cancer cell lines, for 25 of 150 patients. VCaP was sensitive to six therapies according to the GDSC; however, these therapies (selisistat, SB52334, UNC0642, Trichostatin A, acetalax, and linsitinib) do not have an established clinical role in mCRPC (Supplementary Figure 4). Nearest neighbor cell lines to patients with metastatic melanoma were frequently sensitive to MEK and RAF inhibitors, including SB590885 (BRAF inhibitor, nearest neighbor for 11 / 110 patients), refametinib (MEK, 10), RAF_9304 (RAF, 8) and dabrafenib (BRAF, 7) (Figure 3c). Among patients with metastatic melanoma that do not harbor *BRAF* p.V600E but do contain a *NRAS* alteration (n = 24), the most common therapies which recurrent nearest neighbors were sensitive to also included RAF_9304 (3 patients), refametinib (3), and SB590885 (3) (Supplementary Figure 5).

### Integrated clinical interpretation in a prospective precision oncology trial

We lastly compared therapeutic strategies nominated by the complete MOAlmanac methodology with those administered to 83 patients in I-PREDICT (NCT02534675), a prospective clinical trial evaluating personalized therapies based on panel sequencing (Foundation Medicine’s FoundationOne)^35^. Citations and relationships between molecular features and clinical action from the study were reviewed and categorized by MOAlmanac evidence levels (Supplementary Table 3). MOAlmanac processed the 524 molecular features reported for I-PREDICT’s 83 patients on a per-patient basis. Therapies administered in the study (41 unique) or highlighted by our method (40) were categorized by therapeutic strategy according to expert review based on shared pathway targets, resulting in a total of 31 unique strategies (Supplementary Table 3). An overlap in recommended therapeutic strategy was observed in 38 (46%) patients (Supplementary Figure 6). For patient therapy pairs highlighted by MOAlmanac based on FDA evidence or clinical guidelines, 67% and 50%, respectively, were involved in a therapeutic strategy administered by the study. Of the 13 patients with a therapy highlighted by MOAlmanac associated with FDA approved or Guideline evidence that were not involved in an overlapping strategy, 5 patients had another therapy which utilized a strategy administered by I-PREDICT and the remaining 8 nominated therapies approved for other disease contexts. For nominations based on weaker evidence categories, the concordance was 18% for preclinical and 50% for inferential (Figure 4a). The most common concordant strategies were ER signaling inhibition, PI3K/AKT/mTOR inhibition, and immunotherapy (9, 9, and 7 patients, respectively). Of strategies that were not shared, I-PREDICT favored VEGF inhibition for patients with *TP53* alterations (20 patients) whereas MOAlmanac frequently highlighted assertions such as PRMT5 inhibition (13 patients) based on a preclinical relationship showing efficacy of EPZ015666 for *CDKN2A/B* deletions (Figure 4b).

**Figure 4.**
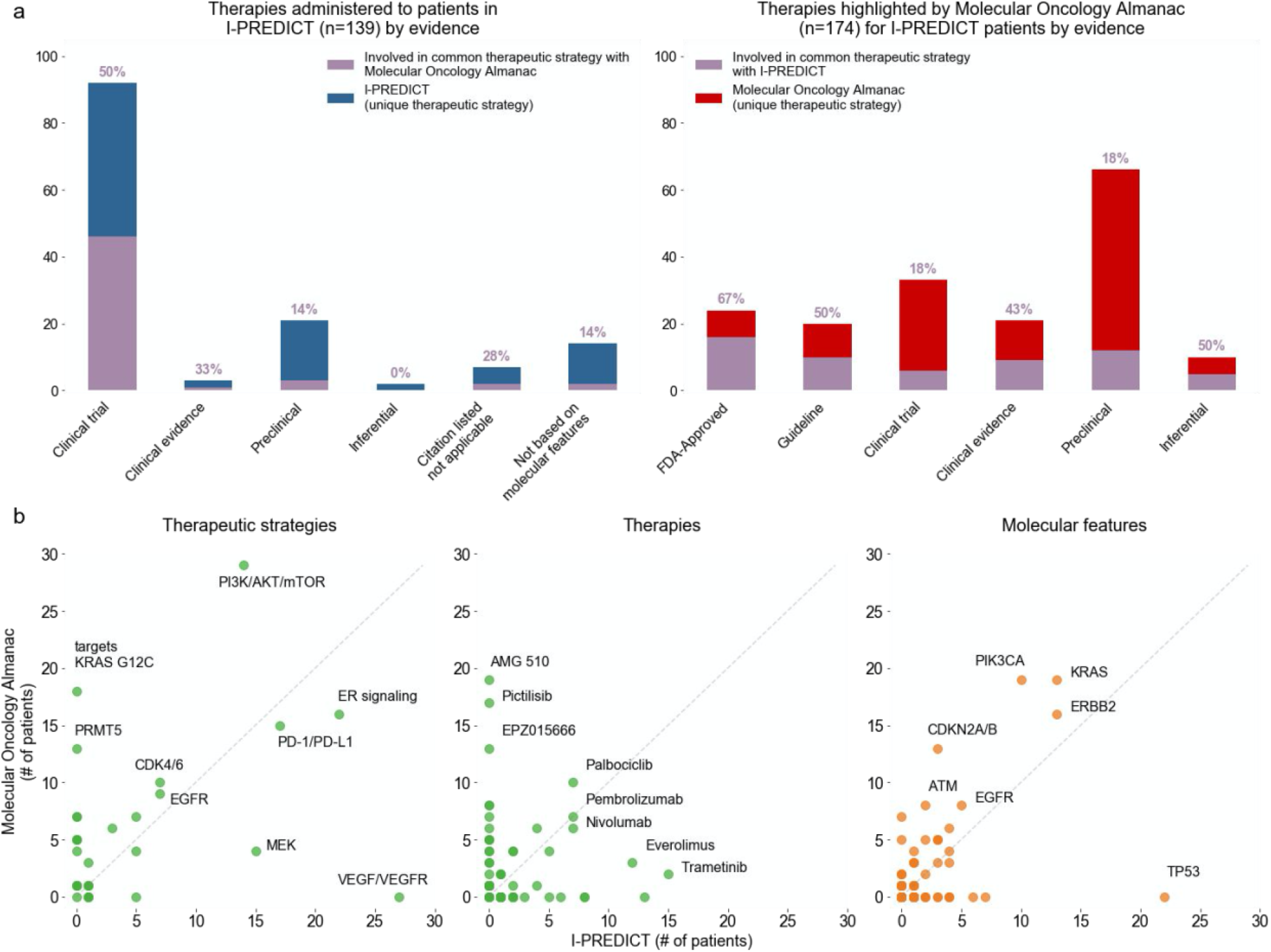
Application of MOAlmanac to a prospective clinical trial. We investigated if MOAlmanac could highlight similar therapeutic strategies that were utilized by real world evidence. MOAlmanac was applied to the I-PREDICT trial, which evaluated the efficacy of molecularly matched therapies in 83 patients^35^. (**a**) Therapies and corresponding molecular features were mapped to therapeutic strategies for those administered in I-PREDICT and highlighted by MOAlmanac. MOAlmanac nominated therapeutic strategies applied for a given patient (purple) more often for those based on well established evidence (i.e. FDA approvals; 67% of therapy patient pairs) relative to less established evidence, such as preclinical (18%). Counts of therapeutic strategies applied to patients that were unique to I-PREDICT are shown in blue and those highlighted by and unique to MOAlmanac are in red. (**b**) Therapeutic strategies, individual therapies, and molecular features as administered or targeted by I-PREDICT and highlighted by Molecular Oncology Almanac.

Finally, using our patient-to-cell line matchmaking module, nearest neighbor cell lines were sensitive to a median of 2 therapies. For example, I-PREDICT administered everolimus and MOAlmanac highlighted AZD8186 and pictilisib in the case of study id 105, a 60 year old female with breast cancer. The nearest neighbor cell line, CAL-29 (bladder carcinoma), was sensitive to taselisib and alpelisib as reported by GDSC2, both of which also target PI3K/Akt/mTOR. In another case, I-PREDICT administered lenvatinib and ramucirumab for VEGF/VEGFR inhibition to study id A009, a 44 year old male with esophageal adenocarcinoma. MOAlmanac highlighted infigratinib for FGFR inhibition for therapeutic sensitivity and the nearest neighbor cancer cell line, A204 (soft tissue), observes sensitivity to both VEGF and FGFR inhibition (VEGF: cediranib, linifanib, motseanib, ponatinib, and tivozanib and FGFR: ponatinib). Thus, MOAlmanac recapitulates established decision making paradigms in a prospective pan-cancer setting and extends potential assertions in new therapeutic directions in other settings.

## DISCUSSION

Here, we present a clinical interpretation method paired with a novel knowledgebase to facilitate decision-making in precision oncology. In addition to first-order feature consideration, MOAlmanac considers second-order molecular features such as mutational signatures, tumor mutational burden, microsatellite stability, and ploidy, as well as high-throughput therapeutic screens of cancer cell lines. Taken together, MOAlmanac addresses two key needs for precision cancer medicine: 1) Point-of-care individualized patient treatment considerations based on complex molecular interactions that considers evidence beyond FDA approvals and clinical guidelines, and 2) Novel therapeutic hypotheses based on integrative interpretations that can be evaluated in preclinical follow up and prospective trials. When applied to retrospective cohorts, we observed that these novel features of MOAlmanac -- assessment of second-order genomic features and consideration of preclinical or inferential evidence -- provided additional hypotheses for prognosis and therapeutic sensitivity and resistance, especially for otherwise variant-negative tumors.

While individual precision oncology studies require fixed versions of alteration-action knowledge bases, rapidly expanding scope of literature on which these databases originate requires constant updating that makes prospective assessment of precision oncology programs difficult. This challenge was evident in comparing MOAlmanac to the I-PREDICT trial, as differences in match selection were driven by differences in therapeutic availability at different time points, variable knowledge capture of the vast precision oncology hypothesis landscape, and levels of evidence to justify treatment selection. These results are suggestive of the urgency to standardize genomic-based clinical trial data and aggregate knowledge bases to parse the vast literature in precision oncology and enable principled, evidence-based clinical care^5,36^. Manual curation of literature is inherently laborious, and prior efforts have encouraged crowdsourcing and meta studies to address this challenge^4,5,37^.

Furthermore, there were areas of note that could specifically improve our evaluation of patient-to-cell line matchmaking for translational hypothesis generation. First, not all cell lines were tested with every therapy; if they were, shared drug response could be characterized in a more nuanced manner than the current boolean status. Second, there is likely an opportunity to develop improved genomic similarity models which align with therapeutic sensitivity. The advent of large, clinically annotated and molecular profiled patient cohorts may enable these techniques and patient similarity networks to be evaluated for precision cancer medicine on patient profiles rather than cancer cell lines^1,38,39^. Indeed, our primary motivation is to develop similarity metrics that account for multiple data types from tumors to properly leverage nearest neighbor approaches. These approaches, which prospectively leverage genomic data rather than retrospectively curated data sources, are imperative to develop therapeutic hypotheses for patients who are variant negative.

In conclusion, MOAlmanac catalyzes the use of expanded feature types, evidence sources, and algorithms for clinical interpretation of integrative molecular features for precision cancer medicine applications. Incorporation of MOAlmanac into future translational studies and clinical trials may directly enable evaluation of the precision oncology hypothesis across patient populations. Furthermore, MOAlmanac can promote evaluation of patient similarity networks using both clinical and preclinical knowledge to aid precision cancer medicine at the individual patient level for translational discovery. The Molecular Oncology Almanac is available at https://moalmanac.org. This method is available on Github (https://github.com/vanallenlab/moalmanac), Docker Hub (https://hub.docker.com/r/vanallenlab/moalmanac), and on the Broad Institute’s Terra (https://portal.firecloud.org/#methods/vanallenlab/moalmanac/). In addition, a web portal to process individual cases through a user interface atop of Terra is available at https://portal.moalmanac.org/. All code related to analyses and figures herein can be found on Github (https://github.com/vanallenlab/moalmanac-paper). Finally, to facilitate crowdsourced updating of MOAlmanac’s knowledge base, Molecular Oncology Almanac Connector (a Google Chrome extension) is available to enable users to nominate relationships with minimal effort.

## METHODS

### Creating a knowledge base

#### Defining a database schema

An SQL schema was planned and abstracted with Vertabelo for cataloging clinical assertions relating molecular features to clinical action. The schema contained four primary abstractions: Assertion, Feature, Source, and Version with additional tables to relate assertion to features and sources; Assertion_To_Feature and Assertion_To_Source, respectively (Supplementary Figure 7). The underlying data structure is implemented as an SQLite database and managed with Python and SQL Alchemy.

The Assertion table is used to catalog a given clinical action. The context of an assertion is catalogued with disease as described in the source (disease), which is mapped to an oncotree code (oncotree_code) and term (oncotree_term), and any applicable disease context such as disease stage (context). If regarding therapeutic sensitivity or therapeutic resistance, the drug name is entered (therapy_name) along with its type (therapy_type: targeted therapy, chemotherapy, radiation, immunotherapy, hormonal therapy, or combination) and a boolean integer of 1 for asserting a relationship to differential therapeutic sensitivity or resistance or 0 for asserting no such relationship. This data structure allows MOAlmanac to capture negative studies documenting that a given feature is not associated with differential therapeutic sensitivity). If regarding prognosis, a boolean integer is entered to suggest a favorable or unfavorable prognosis (favorable_prognosis). The evidence of an assertion is recorded (predictive_implication); available values are “FDA-approved”, “Guideline” for clinical guideline, “Clinical trial” for associations reported from clinical trials, “Clinical evidence” for retrospective studies or human studies not directly reported from a clinical trial, “Preclinical evidence” for findings from mouse models or cancer cell lines, or “Inferential evidence” for findings from mathematical models or an association between molecular features. In some cases, we denote favored assertions (preferred_assertion) to “tie break” otherwise equal assertions based on published literature and clinical use; e.g. Dabrafenib and Trametinib over Vemurafenib for *BRAF* p.V600E. A free text description of the clinical assertion is curated for all entries (description) along with an entry date (created_on) and last modified date (last_updated).

Molecular features are associated with assertions and are catalogued in a flexible manner to accommodate different attributes of a feature type using feature definitions. For example, rearrangements are defined as having a rearrangement type (translocation, fusion), participating genes (gene1, gene2), and a locus; separately, copy number alterations are defined as having a gene, direction, and cytoband. Rearrangements, somatic variants, germline variants, copy number alterations, microsatellite stability, mutational signatures, mutational burden, neoantigen burden, knockdown, silencing, and aneuploidy are currently catalogued with feature definitions. New feature definitions may be easily programmatically defined, allowing the rapid addition of new features without having to modify the underlying data schema.

Sources are catalogued such that all sources will be associated with a citation, source type (abstract, FDA, guideline, journal), and url. Journal articles are further annotated with the associated PubMed ID (PMID) and DOI. Sources regarding a clinical trial will catalog the National Clinical Trial (NCT) registry number.

Version is an unconnected table used to catalog major, minor, and patch numbers of the database.

#### Iterating from TARGET

TARGET catalogued clinical assertions primarily by gene associated with types of recurrent alterations and examples of therapeutic agents paired with an aggregate rationale for the gene. Literature review was performed by curators to review FDA approvals, clinical guidelines, and journal articles to associate clinical assertions from TARGET with a citation. Associations to 52 genes were removed due to insufficient evidence, recent evidence conflicted with the underlying assertion for 1 gene, and 5 genes were partially retained. Ten genes were not migrated to MOAlmanac because we chose to not catalog the underlying assertion type; specifically, we intentionally chose to not include diagnostic relationships and we reclassified biallelic loss to copy number deletions.

#### Cataloging additional assertions

Subsequent curation efforts cataloged FDA approvals, clinical guidelines, conference abstracts, or recently published literature. Relationships were further categorized by the clinical implication of the assertion (therapeutic sensitivity or resistance or prognostic value), therapy type if relevant, and evidence. Genomic feature types considered were somatic and germline variants, copy number alterations, rearrangements, mutational burden, COSMIC mutational signatures, microsatellite stability status, and aneuploidy.

The knowledge base contained 722 assertions which relate molecular features to therapeutic response and prognosis and 4 related to adverse event risk, manually curated from literature review of FDA approvals (87 assertions), clinical guidelines (187), published journal articles (446), and abstracts (5). In addition to characterizing targeted therapies (417 relationships), we have catalogued relationships related to immunotherapies (48), chemotherapies (40), radiation (19), hormonal treatments (7), and combination therapies (11, Figure 1c).

No further assertions were added to MOAlmanac past March 23rd, 2020 for the purposes of this study.

#### Comparison to other knowledge bases

Molecular Oncology Almanac was categorically compared to CIViC and OncoKB, two similar precision oncology knowledge bases, across the categories of therapy types, molecular feature types, assertion types, catalogued evidence, curation type, accessibility, number of assertions, and counted therapy types (Supplementary Table 4). Citations with PubMed reference numbers (PMIDs, 458 citations) were compared and we observed similar findings to previous meta-studies, that no one database subsumes another (Supplementary Figure 8)^37^.

### Developing a clinical interpretation method

#### Accepted inputs

Molecular Oncology Almanac accepts any combination of somatic variants, copy number alterations, rearrangements, germline variants, somatic variants from another source such as a validation sequencing, and breadth of coverage. In addition, several single value or boolean features are passable such as the purity and ploidy of the tumor as float values, a categorical input for microsatellite stability status, a boolean flag to note whole genome doubling. Free text fields are also available to enter a patient or sample id, tumor type, stage, and general description of the molecular profile.

Input files to MOAlmanac have expectations on their format, which can be found on the method’s Github. Somatic, both primary or validation sequencing, and germline variants conform to the National Cancer Institute’s Genomic Data Commons MAF v1.0.0 format, requiring: Hugo_Symbol, Chromosome, Start_position, End_position, Reference_Allele, Tumor_Seq_Allele1, Tumor_Seq_Allele2, Variant_Classification, Protein_Change, Tumor_Sample_Barcode, Normal_Sample_Barcode, t_ref_count, t_alt_count. MOAlmanac is coded to accept input columns based on Oncotator for these inputs; however, this can be changed by editing the colnames.ini file^40^. MOAlmanac currently is coded to accept total copy number alterations produced by ReCapSeg, or GATK3 CNV, and annotated by Oncotator, requiring the columns gene, segment_contig, segment_start, segment_end, sample, and segment_mean^41^. MOAlmanac is coded to accept rearrangements directly from STAR Fusion, requiring the columns fusion_name, SpanningFrags, LeftBreakPoint, and RightBreakPoint^42^. Breadth of coverage is the sum of calculable bases used to call somatic variants, and is required to calculate nonsynonymous mutational burden; a text file containing the integer, such as summing MuTect 1.0’s call stats output, suffices.

The input arguments stage, purity, ploidy, and description are only used for display as metadata in the produced actionability report. Provided tumor types are mapped to standardized ontology terms and codes using Oncotree (http://oncotree.mskcc.org/#/home), if possible. Patient ID is also used as metadata and is also used as a prefix to label all generated outputs.

#### Annotation and evaluation of individual molecular features

Somatic variants, copy number alterations, and gene fusions are annotated with MOAlmanac, Cancer Hotspots, 3D Hotspots, Cancer Gene Census (CGC), Molecular Signatures Database (MSigDB), and COSMIC ^18,19,21–23^. Genomic events are first annotated for their gene presence (1 for present, 0 for wild type) and then receives a higher integer score if applicable; for example, somatic variants whose protein change appears in Cancer Hotspots will be noted by a 2.

Somatic and germline variants are also annotated with ClinVar and ExAC to identify pathogenic or likely pathogenic variants and common variants ^24,25^. Somatic variants and copy number alterations are annotated and evaluated based on a heuristic similar to PHIAL, sorting to events based on their presence in data sources (Supplementary Figure 1a).

MOAlmanac considers individual non-synonymous variants (missense, nonsense, nonstop, frameshift, insertions, and deletions), copy number alterations that are outside of 1.96 standard deviations from the mean of unique segment means (above 97.5 percentile for amplifications and below 2.5 percentile for deletions), and at least 5 spanning fragments for fusions. Events which meet these criteria will be scored by MOAlmanac’s somatic heuristic and be provided in the output file with the suffix “.somatic.scored.txt”, while filtered alterations are made available in the output noted by the “.somatic.filtered.txt” suffix.

For genomic alterations whose gene appears in Molecular Oncology Almanac, the clinical relevance will be labeled based on the match to the catalogued molecular feature and evidence tier of the matched relationship. Complete matches to explicit features (e.g. protein change for variants, direction for copy number alteration, or fusion and partner) will be labeled as Putatively Actionable whereas partial matches or incompletely characterized features (the gene is catalogued of that data type; e.g. a *ETV6-NTRK1* fusion matches to an assertion of *NTRK1* fusions) is labeled as Investigate Actionability. If an alteration’s gene appears in Molecular Oncology Almanac but not catalogued as the same data type, the alteration will be labeled as Biologically Relevant and is not associated with any clinical relationships. For each provided genomic feature, a match is searched for relationships associated with therapeutic sensitivity, resistance, and disease prognosis and, if either labeled as Putatively Actionable or Investigate Actionability, evidence level of the association, therapy name and therapy type (if sensitivity or resistance) or favorable prognosis, relationship description, citation, and URL for the citation are associated. These actionable features are made available in the output file with the suffix “.actionable.txt”.

In addition, a few outputs regarding germline variants are highlighted and made available, if provided (Supplementary Figure 1b). Variants in genes related to hereditary cancers, based on a panel of 83 genes commonly used for germline testing, are produced in an output with the suffix “.germline.hereditary_cancers.txt”^43^. Likewise, variants in genes noted by the American College of Medical Genetics and Genomics secondary findings v2 ^44^ are highlighted in the output with the suffix “.germline.acmg.txt”. Lastly, germline variants in genes related to somatic cancers (based on a gene presence in MOAlmanac, Cancer Hotspots, or Cancer Gene Census) are noted in the output of the suffix “.germline.cancer_related.txt”. Germline variants which match to MOAlmanac will also be included in the actionable output if (1) they are not labeled as common in ExAC (an allele frequency greater than 1 in 1,000 alleles), (2) are labeled as a pathogenic or likely pathogenic variant in ClinVar, or (3) a truncating (frameshift, nonsense, nonstop, or splice site) variant.

If somatic single nucleotide variants are provided for both primary and secondary (also referred to as validation or orthogonal sequencing) sequencing, MOAlmanac will annotate variants called in the primary sequencing based on their presence (allelic fraction and coverage) in the secondary sequencing. The power to detect variants in the secondary sequencing is calculated using a beta-binomial distribution with *k* equal to 3 for a minimum of three reads, *n* as coverage of the variant in secondary sequencing, *alpha* and *beta* defined as the alternate and reference read counts + 1 as observed from the primary sequencing, respectively. This approach is consistent with best practices by Yizhak et al. 2019 with RNA MuTect^11^. The allelic fraction of somatic variants observed in primary and orthogonal sequencing are plotted against each other in a scatter plot in the output of the suffix “.validation_overlap.png”, with variants observed with detection power greater than or equal to the specified minimum (default 0.80) colored in blue and those otherwise grey. At the moment, MOAlmanac only leverages orthogonal sequencing for validation and does not use it for discovery. When applied to the retrospective cohorts of metastatic melanoma and mCRPC, we had sufficient power to observe 190 of 453 applicable clinically relevant variants. Of note, *AR* p.L702H and p.T878A, variants putatively associated with resistance to androgen deprivation, were observed in the RNA of 6 and 4 patients, respectively^45^.

#### Annotation and evaluation of integrative and second-order genomic features

To ease the process of reviewing multiple intra-gene alterations, MOAlmanac summarizes all somatic variants, germline variants, copy number alterations, and fusion events per gene for genes found within MOAlmanac, Cancer Hotspots, and Cancer Gene Census. Any genes with at least one alteration across any data type will be reported in the output with the suffix “.integrated.summary.txt”.

Somatic alterations are annotated with the number of frameshift, nonstop, nonsense, or splice site germline events within the same gene. This count is labeled as the column “number_germline_mutations_in_gene” in the output of the suffix “.somatic.scored.txt”.

Tumor mutational burden (TMB) is calculated based on the number of nonsynonymous variants divided by the somatic calculable bases. TMB is compared to values calculated for TCGA molecular profiles by Lawrence et al. 2013 to yield a pancan percentile and tissue-specific percentile, if ontology matched to one of the 27 tumor types studied in the publication^46^. TMB for a molecular profile is designated as high if greater than 10 nonsynonymous variants per megabase and greater than or equal to the 80th tissue-specific percentile, or pancan percentile if not mapped.

COSMIC mutational signatures are evaluated using deconstructSigs by running R as a subprocess using the default trinucleotide counts method ^47,48^. Signatures with a contribution greater than a specified minimum contribution (default: 0.20) are annotated at least as Biologically Relevant and annotated using MOAlmanac for consideration of actionability. Nucleotide context counts are made available in table format directly from deconstructSigs as an output with the suffix “.sigs.context.txt” and signature contributions with the suffix “.sigs.cosmic.txt”. Trinucleotide counts of a considered molecular profile are plotted based on raw and normalized counts in the outputs “.sigs.tricontext.counts.png” and “.sigs.tricontext.normalized.png”, respectively.

Microsatellite stability is both directly considered as a categorical input for status and indirectly by highlighting potentially related variants. As a direct input, users may flag microsatellite status as microsatellite stable, microsatellite instability low, microsatellite instability high, or unknown. Genomic alterations which appear in genes related to microsatellite instability are highlighted as supporting variants and Biologically Relevant and further noted in their own output, with the suffix “.msi_variants.txt”; specifically, the genes considered are *ACVR2A, DOCK3, ESRP1, JAK1, MLH1, MSH2, MSH3, MSH6, PMS2, POLE, POLE2, PRMD2*, and *RNF43* ^49,50^. As of this publication, MOAlmanac has only catalogued assertions related to MSI-High status.

Whole genome doubling, or aneuploidy, is available for consideration as a boolean-valued input and, if flagged, will evaluate for clinical relevance based on the currently catalogued assertions. As of this publication, MOAlmanac has catalogued Bielski et al. 2018’s observation that whole genome doubling being associated with adverse survival across a pan-cancer setting^28^.

Mutational burden, mutational signatures, microsatellite stability, and whole genome doubling are at most highlighted as Investigate Actionability by Molecular Oncology Almanac for clinical assessment.

#### Creating clinical actionability reports

Clinical actionability reports are created for all profiles processed with Molecular Oncology Almanac, generated with Python 3.6, Flask, and Frozen Flask.

The reports contain sections containing profile metadata (Profile Information), molecular features associated as a Putatively Actionable or Investigate Actionability predictive implication for therapeutic sensitivity or resistance and prognosis, as well as variants associated with Biological Relevance (Actionability Report). Associations list the implication, evidence, and associated therapy and description of clinical assertion as rationale. Sources for each association are available as hyperlinks labeled as “[source]”, equivalent assertions are available to view in a modal labeled, and preclinical efficacy of the assertion is also available as modal, if applicable.

The 5 most similar cell lines to the provided molecular profile are listed by their CCLE name along with their sensitive therapies and clinically relevant features. For each cell line, a modal is available that lists their Broad/DepMap and Sanger Institute aliases and somatic variants, copy number alterations, and fusions in any MOAlmanac, Cancer Hotspot, or CGC gene as well as the ln(ic50), AUC, and z score for each of the top 10 most sensitive therapies of the cell lines. This feature can be hidden in the clinical report passing diable_matchmaking as a parameter to the method.

Due to being produced with Frozen Flask, these web based reports are a single html file with no additional file dependencies. They usually are no larger than 1 Mb in size.

### Comparing PHIAL-TARGET and MOAlmanac with two retrospective studies

#### Data acquisition and sample processing

Whole-exome sequencing (WES) and RNA sequencing (RNA-seq) was acquired for 110 previously published patients with metastatic melanomas (n = 44 with RNA)^26^ and 150 patients with castration-resistant prostate cancers (mCRPC, n = 149 with RNA)^27^. Subsequent sample processing was performed on the Broad Institute and Verily Life Sciences’ Terra Google Cloud platform.

Whole-exome sequencing was used to call somatic and germline variants and copy number alterations. MuTect 1.0 was used to identify single nucleotide variants (SNVs) and somatic calculable bases of individual tumor samples while Strelka was used to identify insertions and deletions (InDels)^51,52^, run utilizing the Getz Lab CGA WES Characterization pipeline at the Broad Institute. Artifacts introduced by DNA oxidation during the sequencing process were removed ^53^. Mutations calls were compared to a panel of germline samples and were removed if they appeared in more than three germline samples^54^. Germline variants were called using Deep Variant^55^. Segmented total copy number was calculated across the exome by comparing fractional exome coverage to a panel of normals using CapSeg ^56,57^. Tumor purity and ploidy was calculated using FACETS^58^.

Transcriptome BAMs were converted to FASTQ format and aligned using STAR ^59^. Fusions were then called using STAR Fusion^60^. STAR aligned bams were calibrated following GATK’s best practices for variant discovery in RNA-seq (https://github.com/broadinstitute/gatk-docs/blob/3333b5aacfd3c48a87b60047395e1febc98c21f9/gatk3-methods-and-algorithms/Calling_variants_in_RNAseq.md) using GATK 3.7^61–63^. Somatic variants observed in whole-exome data were then force called from the recalibrated RNA-seq bams for each individual using MuTect 1.0.

Somatic variants from both WES and RNA-seq, germline variants, and copy number alterations were annotated using Oncotator v1.9.1^40^.

#### Comparison of clinically relevant events

Molecular features were processed for all 260 samples by both PHIAL 1.0.0 (https://github.com/vanallenlab/phial)^2^ and MOAlmanac. While both methodologies considered all available genomic events, PHIAL considered somatic variants and copy number alterations while MOAlmanac additionally considered germline variants, rearrangements, mutational burden, mutational signatures, and whole-genome doubling. Microsatellite stability was not considered for this analysis as labels from testing, if performed, were not available. Events that matched with the underlying knowledge base as either Investigate Actionability or Putatively Actionable, thus stronger than simply a gene match, were considered for clinical relevance (Supplementary Figure 2). While the differences were impacted by literature curation and MOAlmanac considering additional feature types, they were also impacted by changing how copy number alterations are handled; PHIAL called copy number alterations based on a threshold, |segment mean| ≥ 1, whereas MOAlmanac utilizes a percentile approach, top or bottom 2.5%.

### Expanded methods for directly leveraging preclinical models

#### Data acquisition and processing

Somatic variants and copy number alterations for cancer cell lines catalogued in the Cancer Cell Line Encyclopedia were gathered from cBioPortal and fusions and therapeutic sensitivity were downloaded from the Sanger Institute’s Genomics of Drug Sensitivity in Cancer (GDSC) ^32,33^. Cancer cell lines were standardized by name and filtered for by requiring: all four data types being available, being of solid tumor origin, not subject to genetic drift between Broad and Sanger versions of the cell line per Ghandi et al. 2019, and not reclassified as fibroblast like by Weck et al. 2017 and Ghandi et al. 2019 ^32,64^; resulting in 452 cancer cell lines. Somatic variants, copy number alterations, and fusions were formatted for usage and annotated by Molecular Oncology Almanac.

#### Directly leveraging preclinical models to evaluate efficacy

All GDSC1 and GDSC2 therapies were mapped to therapies catalogued in MOAlmanac. For all therapies associated with genomic events by MOAlmanac for which a GDSC mapping exists, a sensitivity dictionary is created in which each key is associated with a clinically relevant feature found by the method. For each feature, we list all mutant and wild type cell lines for each component; e.g. when considering *CDKN2A* deletions, mutant and wild type lists are made for all cell lines that have any alteration in *CDKN2A* (somatic variant, copy number alteration, or fusion), cell lines that have a *CDKN2A* copy number alteration, and cell lines that have a *CDKN2A* deletion. For each pairing of mutant and wild type cell lines, the IC50 values are compared with a Mann-Whitney-Wilcoxon test to evaluate if a significant difference exists between the two distributions. A box plot of mutant and wild type cell lines and their IC50 values is also created, labeled by the genomic feature used to stratify.

The results of such testing are reported in two outputs, the actionability report with the suffix “.report.html” and a table compiling all examinations with the suffix “.preclinical.efficacy.txt”. When applicable, a hyperlink labeled as “[Preclinical evidence]” will appear under “Therapy & rationale” for variants and features associated with therapeutic sensitivity. Upon clicking the link, a modal window opens showing all box plots of comparisons along with the number of wild type cell lines, number of mutant cell lines, and the Mann-Whitney-Wilcoxon statistic and p-value for each feature evaluated. In addition, IC50 median, mean, and standard deviation can be found for all relationships evaluated in the mentioned preclinical efficacy table output.

#### Directly leveraging preclinical models for patient-model matchmaking

We sought to directly leverage molecular profiles for clinical interpretation. For the purposes of this application, we sought to compare a case molecular profile to a larger population and sort other members by genomic features such that the nearest neighbor to our case profile shared drug sensitivity. In absence of a large cohort of clinically annotated primary or metastatic tumor profiles, we utilized cancer cell lines which have been characterized by high throughput drug screens and evaluated by comparing cell lines against cell lines.

GDSC z scores of therapies applied to cell lines were utilized to convert continuous valued IC50 response curves to boolean valued sensitive (z score ≤ −2) or resistant (z score ≥ 2)^33^. Pairwise comparisons were made between all cell lines which contained GDSC therapeutic response data, noting the intersection of therapies which both profiles were deemed sensitive to as well as the intersection size. If the intersection size was greater than 0, the pair was deemed to share therapeutic sensitivity. When evaluating a novel case profile the matchmaking module of MOAlmanac, the 452 cancer cell lines, which result from filtering described in two sections prior, are used for comparison. However, for evaluation, we further required that cell lines are sensitive to at least one therapy and that there exists at least one other cell line that shares therapeutic sensitivity, so that there is at least one true positive when sorting other cell lines, resulting in 377 cell lines.

After somatic variants, copy number alterations, and fusions were annotated and evaluated by MOAlmanac, molecular features were vectorized into sample x feature tables. The coding of features was dependent on the model implemented, discussed more explicitly in the next section; however, some commonalities exist. All elements were boolean valued and thus all feature tables were sparse boolean arrays. When a similarity model involved genes, either the CGC (n = 719 genes) or MOAlmanac (130) gene sets were used. Among a series of models tested, we found the best performing model to be using Similarity Network Fusion on four sample x feature tables: CGC genes altered by somatic variants, copy number alterations, and fusions and a fourth table of samples x specific molecular features associated with an FDA approved therapy, subsequently referred to as SNF: CGC & FDA.

##### Evaluation metrics were borrowed from ranked retrieval

The performance of how a similarity metric sorts cell lines relative to one cell line are evaluated using precision @ rank (*k*), recall @ *k*, and average precision. Consider four cell lines sorted in order relative to a case profile such that the first and third share therapeutic sensitivity with the case profile and the second and fourth does not (Figure 3a). Cell lines which share therapeutic sensitivity can be considered relevant. To calculate precision @ *k*, given *k* neighbors, we divide the number of relevant neighbors divided by *k*; e.g. considering the first neighbor (k=1) yields a precision @ 1 of 1.0 (1 relevant neighbor / 1) but considering the second neighbor as well yields a precision @ 2 of 0.5 (1 relevant neighbor / 2). Recall is calculated as the fraction of overall relevant neighbors returned when considering *k* neighbors; at *k = 1* recall is calculated to be 0.5 in our example, until *k* = 3 when a second relevant cell line is returned thus recall is calculated to be 1.0, and recall = 1.0 at *k* = 4. Average precision (AP) is calculated by taking the average of precision values at positions of a relevant neighbor; using our example, relevant neighbors exist at precision @ k = 1 and 3 with associated precision values of 1.0 and 0.66 so the average precision for this sort, or query to use terminology from information retrieval, is calculated to be 0.83.

The performance of a similarity metric for many queries can be evaluated by calculating the mean average precision (mAP). Given three case profiles which sorted cell lines against them with average precision values of 0.66, 0.565, and 0.25, the mean average precision is the average of them, which is calculated to be 0.492. In our context, for each similarity model, we calculate the average precision for each cell line and the mean average precision across all cell lines (Supplementary Table 5).

Models can be compared pairwise with permutation testing (Supplementary Table 6). The difference in mean average precision (delta mAP) is chosen as a test statistic and the AP @ k values are shuffled for all 377 values of k. Given these shuffled AP @ k values, mAP values are calculated along with a delta mAP and the delta mAP is recorded. This was performed over 10,000 iterations using seeds 0 to 9,999 to create a distribution of delta mAP values. The test statistic is compared to the distribution to generate a p-value and, if the p-value was ≥ 0.05, it was deemed that the two models were within the noise range of one another. Our best performing model SNF: CGC & FDA was within the noise range of two other models, a multi-pass sort of first using agreement based measure of molecular features associated with an FDA approved therapy followed by agreement based sort of CGC genes mutated by any feature type (Multi-pass sort: FDA & CGC, p=0.4013) and sorting cell lines by their mutant and wild type status of variants in order based on the somatic heuristic in MOAlmanac (Somatic tree, p=0.5458); however, SNF: CGC & FDA observed a stronger AP @ k = 1 in both cases, 0.193 versus 0.164 and 0.119, respectively.

There are several areas which we note that this framework could be improved. First, not all cell lines were treated with all therapies and we can not deem an untested pair as sensitive or not sensitive unless we resort to estimating missing data, thus, we assume that cell lines do not respond to therapies which they were not tested to be conservative in our analysis. In the setting of a complete pairing (all cell lines are treated with all therapies) we could incorporate a more nuanced label. For example, we could continue using the z score thresholds but instead label based on the jaccard index of shared therapies or we could transition to using a continuous valued similarity of drug sensitivity such as euclidean distance of IC50s or perform a PCA. In either case, a complete pairing of therapies and cell lines would enable us to use additional evaluation metrics such as Discounted Cumulative Gain (DCG), ranking other cell lines based on a relevance scale rather than a boolean condition and rank. Secondly, rather than evaluate cell lines against cell lines, we envision that an ideal experiment for this analysis would involve a cohort of paired primary tumor samples and patient derived cell lines in which we would hope that the paired patient derived cell line would be deemed most similar to its corresponding tissue sample. Such a setting would enable the studying of performance as a function of cell line passages. Expression was not used in this analysis as it is a feature modality not yet commonly used at the point-of-care.

#### Models and calculating similarity metrics

Several models were implemented to characterize similarity between cancer cell lines based on genomic features. Models were evaluated using average precision, specifically average precision @ k = 1, and mean average precision. In short, our best performing model (SNF: FDA & CGC) observed a AP @ k = 1 of 0.194 which was 2.03x better than random but still only recommends a nearest neighbor for one fifth of cell lines. We are excited to see improvements in directly leveraging molecular profiles for clinical interpretation. Performance of models can be found in Supplementary Table 5. Models include, listed alphabetically:

Compatibility (compatibility). Inspired by dating algorithms, we weigh each molecular feature (or question) based on strength of the match (e.g. a BRAF deletion only matches BRAF p.V600E by gene). With these relative weights, we calculate a max score for each sample and compare against other cell lines.

Jaccard of MOAlmanac feature types (jaccard-almanac-feature-types). We sort by agreement based measure (jaccard) by considering both gene and data type for all somatic variants, copy number alterations, and rearrangements catalogued in the Molecular Oncology Almanac (e.g. CDKN2A copy number alterations match but not a CDKN2A deletion and CDKN2A nonsense somatic variant).

Jaccard of MOAlmanac features (jaccard-almanac-features). We sort by agreement based measure (jaccard) by considering all somatic variant, copy number, and rearrangement molecular features catalogued in the Molecular Oncology Almanac.

Jaccard of MOAlmanac genes (jaccard-almanac-genes). We sort by agreement based measure (jaccard) by considering any somatic variant, copy number alteration, and rearrangement in any gene catalogued in Molecular Oncology Almanac.

Jaccard of CGC feature types (jaccard-cgc-feature-types). We sort by agreement based measure (jaccard) by considering variants in a Cancer Gene Census gene and feature type (e.g. CDKN2A copy number alterations match but not a CDKN2A deletion and CDKN2A nonsense somatic variant).

Jaccard of CGC genes (jaccard-cgc-genes). We sort by agreement based measure (jaccard) by considering any variant in a Cancer Gene Census gene.

Multi-pass sort: FDA & CGC (multi-pass-sort_fda-cgc). A weakness of agreement based measure is that there will be tied values. We tie break similarities based on Molecular Oncology Almanac features associated with FDA evidence by using similarity based on CGC genes.

Nonsynonymous variant count (nonsynonymous-variant-count). We assign neighbors based on the absolute value of the difference of the number of coding somatic variants. This is a proxy for mutational burden, because we do not have the number of somatic bases considered when calling variants to use a denominator.

PCA of MOAlmanac genes (pca-almanac-genes). We run PCA and then nearest neighbors for the vectorization of MOAlmanac genes, with mutants being without consideration of feature type. For example, there is one feature called “TP53” and both TP53 nonsense variants and copy number deletions can populate the element.

PCA of CGC genes (pca-cgc-genes). We run PCA and then nearest neighbors for the vectorization of CGC genes, with mutants being without consideration of feature type. For example, there is one feature called “TP53” and both TP53 nonsense variants and copy number deletions can populate the element.

Random (random_mean). Randomly shuffle cell lines against one another across 100,000 seeds. This uses the seed of the average mean average precision.

SNF: MOAlmanac (snf_almanac). Rather than collapse all data types into a single similarity matrix (e.g. with columns such as CDKN2A somatic variant, CDKN2A copy number alteration), we use the python implementation of Similarity Network Fusion by Ross Markello(https://github.com/rmarkello/snfpy)^34^. We fuse networks that describe agreement based on variants in almanac genes in (1) somatic variants, (2) copy number alterations, and (3) rearrangements.

SNF: CGC (snf_cgc). Rather than collapse all data types into a single similarity matrix (e.g. with columns such as CDKN2A somatic variant, CDKN2A copy number alteration), we use the python implementation of Similarity Network Fusion by Ross Markello (https://github.com/rmarkello/snfpy)^34^. We fuse networks that describe agreement based on variants in CGC genes in (1) somatic variants, (2) copy number alterations, and (3) rearrangements.

SNF: FDA & CGC (snf_fda-cgc). We perform similarity network fusion using the python implementation by Ross Markello (https://github.com/rmarkello/snfpy) to fuse networks that contain: (1) CGC genes that contain a somatic variant, (2) CGC genes that contain a copy number alteration, (3) CGC genes that contain a rearrangement, (4) Almanac features associated with FDA evidence^34^.

SNF: FDA & CGC genes (snf_fda-cgc-genes). We perform similarity network fusion using the python implementation by Ross Markello (https://github.com/rmarkello/snfpy) to fuse networks that contain (1) almanac features associated with FDA evidence and (2) any variant occurring in a Cancer Gene Census gene.

Somatic tree (somatic-tree). This is somewhat inspired by CELLector by Najgebauer et al.^16^. One issue with agreement based measures is that each feature is weighted the same. CELLector has a sorted list of genes/variants based on cancer type and will report similar cell lines based on mutant / wild type status of each gene. While not exactly the same, we use the annotations from various data sources appended to variants by Molecular Oncology Almanac to create a priority list for variants (hotspots ranked the highest, etc.). For each case sample, we consider the genes which are observed to be mutated and preserve the order that they would appear in the somatic.scored.txt output of MOAlmanac. All other samples are then sorted by their mutant / wild type status of these genes.

### Comparing to a prospective clinical trial, I-PREDICT

We compared the clinical actions administered based on molecular profiles to patients in the I-PREDICT prospective clinical trial to those highlighted by Molecular Oncology Almanac^35^. All genomic events considered were present in the supplementary text of the study and we extracted molecular features, therapies administered, and citations. Disease ontologies were mapped to Oncotree terms and codes (http://oncotree.mskcc.org/). Molecular features were formatted for annotation and evaluation by MOAlmanac.

Citations providing rationale for therapies administered based on molecular features were extracted from the supplementary text, obtained, read, commented on, and categorized by evidence level. Molecular features considered by the study were merged with annotations made by MOAlmanac and, using the author notes from the supplementary text, we annotated if the study targeted the molecular feature. Therapy and associated molecular features were mapped to therapeutic strategies by expert review. Therapies administered in the study and those highlighted by MOAlmanac for therapeutic sensitivity were listed on a per patient basis and evidence levels were annotated for each therapy per patient. For therapies administered by the study, citations cited per patient were referenced again for the specific relationship between therapeutic strategy or therapy and molecular feature. Each therapy administered was binned based on the evidence level or annotation as no citation, if the therapy was administered not on the basis of molecular features, or citation listed not applicable, if the citation(s) listed did not mention the therapy, strategy, or target. In some cases which would have resulted in the latter, we transcribed that perhaps a source cited for another relationship in the cohort and cited that source. Therapies were tagged with a boolean value if they were involved in a shared therapeutic strategy between what was administered in I-PREDICT and highlighted by Molecular Oncology Almanac for a given patient (Supplementary Table 3).

### Web-based tools to improve accessibility

#### Browsing the knowledge base

A web based browser was created for browsing the knowledge base with Python, Flask, and SQLAlchemy and hosted on Google Compute Engine, herein referred to Molecular Oncology Almanac Browser or browser. The front page lists the total number of molecular features and assertions catalogued as well as the total number of cancer types, evidence levels, and therapies entered. A central search box allows for searching across multiple search terms such as evidence, gene, feature types, or feature type attributes (protein changes, genomic positions, etc.). The browser also features an about page, which contains a hyperlink to download the contents of the knowledge base. Users may submit entries for consideration into the database with a web form, accessible through the “Submit entry” menu item.

#### Application Program Interface (API)

To interact with the knowledge base programmatically, an application program interface (API) was built using Python and Flask to interface with the browser’s underlying data structure. Several get requests are available to list therapies, evidence levels, or genes as well as the ability to get all or by id assertions, sources, feature definitions, features, feature attribute definitions, or feature attributes. A post request is available to suggest a new assertion to the database.

#### Reducing the burden of crowdsourcing

To reduce the burden of crowdsourcing, we created a Google Chrome extension, herein referred to as Molecular Oncology Almanac Connector or connector, with Python and Flask. The connector allows users to submit a DOI along with a feature type, cancer type, evidence level, and therapy if relevant. The user’s email address is also requested in order to follow up about the nominated assertion. This is accomplished using the post request API endpoint for new assertions. The privacy policy of the Connector was reviewed and approved by Dana-Farber compliance.

#### Creating a cloud-based execution portal

A web portal was built using Python, Flask, and requests to take advantage of Terra’s (formerly known as FireCloud) API and Google Cloud’s gsutil in order to allow run MOAlmanac without needing to use Python, Github, Docker, or Terra. Users must have billing set up with and be registered on Terra and, upon selecting to begin a new analysis, users will be asked to specify a de-identified sample name, either a free text tumor type or select one based on a drop down menu containing ontologies from Oncotree, and a Terra billing project. A workspace will be created in the specified billing project named based on the sample name, tumor type, and a timestamp. The remaining fields are optional and any combination of them can be provided.

Somatic single nucleotide variants, insertions and deletions, bases covered, copy number alterations, fusions, and somatic variants from orthogonal sequencing as well as a free text description can be uploaded to the workspace through the web portal. The privacy policy and application were reviewed and approved by Dana-Farber compliance and information security; Nonetheless, we decided to remove germline inputs via the portal.

Upon submission, a Terra workspace and corresponding Google bucket is created that only the user has access to and provided files are uploaded to the Google bucket. The workspace and data model are populated based on inputs and a submission of Molecular Oncology Almanac is run. The workspace is tagged with the tag Molecular-Oncology-Almanac-Portal on Terra. The user is returned to their homepage on the portal, showing a summary of workspaces submitted through the portal, by subsetting workspaces that they have access for the portal’s tag. The summary will note the job submission until the page. Upon page refresh with the job being completed, a direct hyperlink to view the report output (View Report) is made available.

#### Analysis and data availability

All analyses and figures referenced herein can be found in and regenerated with the paper’s Github repository: https://github.com/brendanreardon/moalmanac-paper. Code is available for all software in the Molecular Oncology Almanac ecosystem: browser (https://github.com/vanallenlab/almanac-browser), connector (Google Chrome extension, https://github.com/vanallenlab/almanac-extension), method (https://github.com/vanallenlab/moalmanac), and portal (https://github.com/vanallenlab/almanac-portal).

## Acknowledgements

We thank Alexander Bauman and Ruchi Munshi of the Broad Institute’s Data Science and Data Engineering Platform for their help with the Terra API as well as Kathleen Tibbits and David Shiga for their mentorship. This work was supported by NIH U01 CA233100, NIH R01 CA227388, NIH R37 CA222574, PCF-Movember Challenge Award, and the Mark Foundation Emerging Leader Award.

## Author information

### Affiliations

**Department of Medical Oncology, Dana-Farber Cancer Institute, Harvard Medical School, Boston, MA, USA**

Brendan Reardon, Nathaniel D Moore, Nicholas Moore, Eric Kofman, Saud Aldubayan, Alexander Cheung, Jake Conway, Haitham Elmarakeby, Tanya Keenan, Daniel Keliher, David Liu, Jihye Park, Natalie Vokes, Felix Dietlein, Eliezer M Van Allen

**Cancer Program, Broad Institute of MIT and Harvard, Cambridge, MA, USA**

Brendan Reardon, Nathaniel D Moore, Nicholas Moore, Eric Kofman, Saud Aldubayan, Alexander Cheung, Jake Conway, Haitham Elmarakeby, Alma Imamovic, Sophia C. Kamran, Tanya Keenan, Daniel Keliher, David J Konieczkowski, David Liu, Kent Mouw, Jihye Park, Natalie Vokes, Felix Dietlein, Eliezer M Van Allen

**Indiana University School of Medicine, Indianapolis, IN, USA**

Nathaniel D Moore

**Howard Hughes Medical Institute, Chevy Chase, MD, USA**

Nathaniel D Moore

**Department of Internal Medicine, University of Cincinnati, Cincinnati, Ohio, USA**

Nathaniel D Moore

**Harvard Medical School, Harvard University, Boston, MA, USA**

Nicholas Moore, Kent Mouw

**Department of Cellular and Molecular Medicine, University of California, San Diego, La Jolla, CA, USA**

Eric Kofman

**Institute for Genomic Medicine, University of California, San Diego, La Jolla, CA, USA**

Eric Kofman

**Division of Genetics, Brigham and Women’s Hospital, Boston, MA, USA**

Saud Aldubayan

**College of Medicine, King Saud bin Abdulaziz University for Health Sciences, Riyadh, Saudi Arabia**

Saud Aldubayan

**Grossman School of Medicine, New York University, New York, NY, USA**

Alexander Cheung

**Division of Medical Sciences, Harvard University, Boston, MA, USA**

Jake Conway

**Department of System and Computer Engineering, Al-Azhar University, Cairo, Egypt**

Haitham Elmarakeby

**Department of Pediatric Oncology, Dana-Farber Cancer Institute, Harvard Medical School, Boston, MA, USA**

Alma Imamovic

**Department of Radiation Oncology, Massachusetts General Hospital, Harvard Medical School, Boston, MA, USA**

Sophia Kamran

**Department of Mathematics, Tufts University, Medford, MA, USA**

Daniel Keliher

**Department of Radiation Oncology, Dana-Farber Cancer Institute & Brigham and Women’s Hospital, Boston, MA**

David J Konieczkowski, Kent Mouw

**Harvard Radiation Oncology Program, Massachusetts General Hospital, Boston, MA, USA**

David J Konieczkowski

**Department of Radiation Oncology, The Ohio State University Comprehensive Cancer Center - Arthur G. James Cancer Hospital and Richard J Solove Research Institute, Columbus, OH, USA**

David J Konieczkowski

**Department of Thoracic / Head and Neck Oncology, MD Anderson Cancer Center, Houston, TX, USA**

Natalie Vokes

### Contributions

Conception and designs: B.R., N.D.M., N.M., E.K., F.D., E.M.V.A. Development of methodology: B.R., N.D.M., N.M., E.K., S.A., A.C., J.C., H.E., A.I., S.C.K., T.K., D.K., D.J.K., D.L., K.W., J.P., N.V., F.D., E.M.V.A. Analysis and interpretation of data: B.R., N.D.M., N.M., E.K., E.M.V.A. Writing, review, and/or revision of the manuscript: B.R., N.D.M., N.M., E.K., S.A., A.C., J.C., H.E., A.I., S.C.K., T.K., D.K., D.J.K., D.L., K.W., J.P., N.V., F.D., E.M.V.A. Study supervision: E.M.V.A.

### Corresponding author

Correspondence to Eliezer M Van Allen

