## Supplementary figures and table captions for "Clinical interpretation of integrative molecular profiles to guide precision cancer medicine"

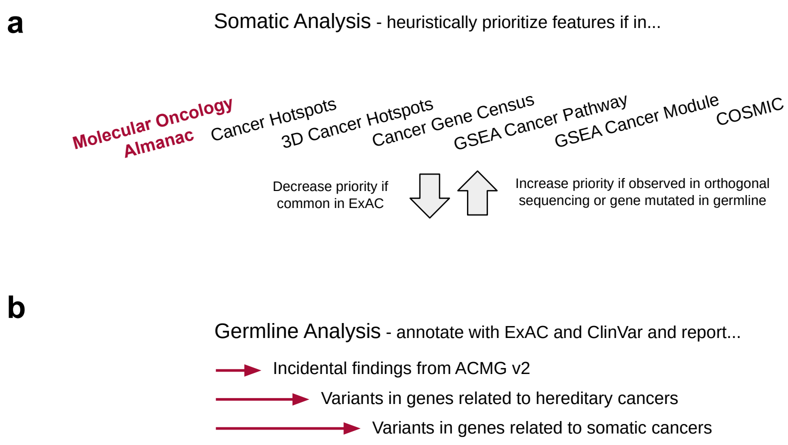


**Supplementary Figure 1.** Schematics of heuristics for sorting of somatic and germline variants.

(**a**) Like PHIAL, the Molecular Oncology Almanac heuristically evaluates somatic variants and will decrease variant priority if likely to be a common variant or increase variant priority if observed in validation sequencing. (**b**) The Molecular Oncology Almanac will annotate provided germline variants for pathogenic or likely pathogenic status of rare or uncommon variants to report incident findings based on the American College of Medical Genetics v2, variants in genes related to hereditary cancers, and variants in genes related to somatic cancers[21,43,44].

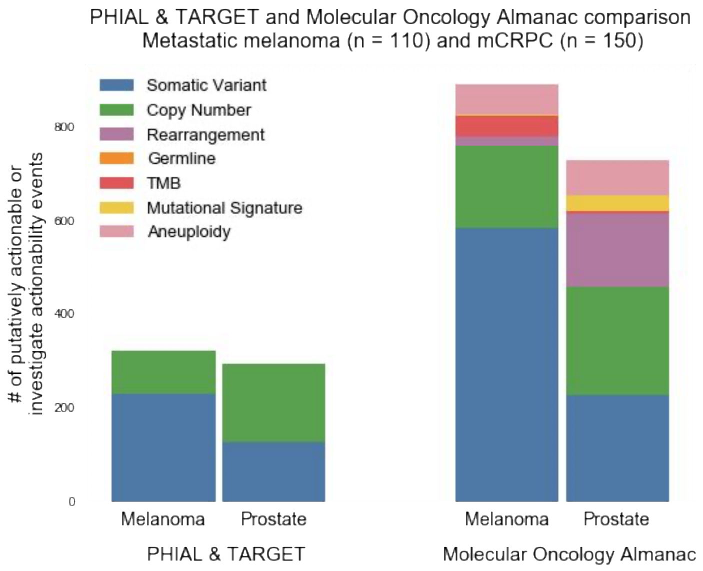


**Supplementary Figure 2.** Clinically relevant molecular features labeled by PHIAL & TARGET vs Molecular Oncology Almanac.

Counts of molecular features labeled as either putatively actionable or investigate actionability, thus being associated with a clinical action, by PHIAL & TARGET vs Molecular Oncology Almanac. Both methodologies were provided with the same list of features and differences resulted from improved handling of genomic features (e.g. percentile approach for copy number instead of thresholded), expanded evidence (MOAlmanac contains more relationships between molecular features and clinical relationships), and expanded feature types (whereas PHIAL & TARGET only included somatic variants and copy number alterations, MOAlmanac contains more). This does not include “Biologically Relevant” features from MOAlmanac or “High Priority” from PHIAL & TARGET, neither of which are directly associated with assertions.

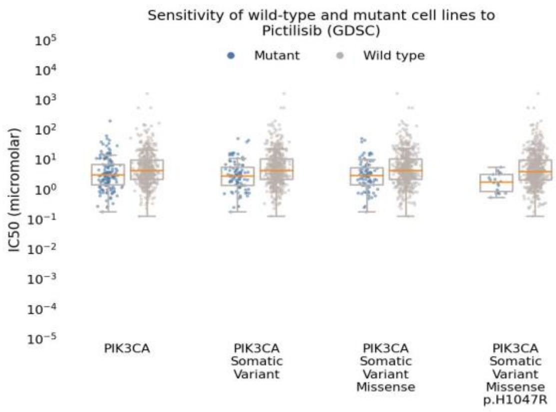


**Supplementary Figure 3.** MOAlmanac inspects preclinical efficacy of highlighted relationships. Sensitivity of Pictilisib on 452 cancer cell lines for all *PIK3CA* mutations, all *PIK3CA* somatic variants, *PIK3CA* missense variants, and *PIK3CA* p.H1047R shown.

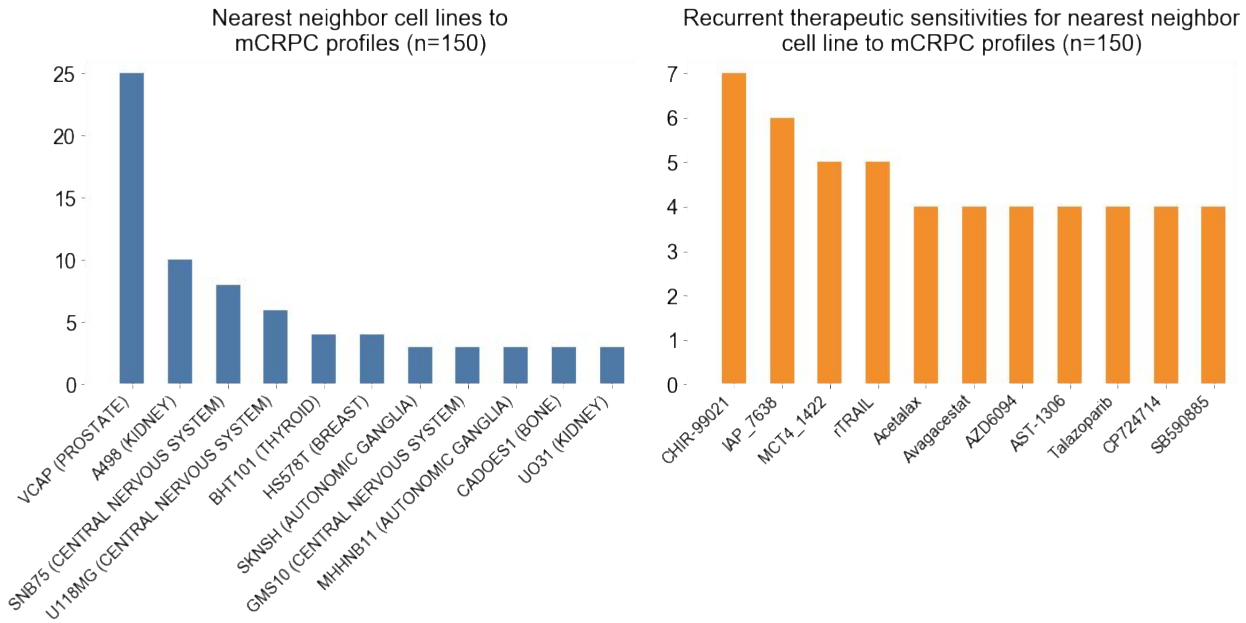


**Supplementary Figure 4.** Patient-to-cell line matchmaking applied to 150 mCRPC profiles. Of 150 patients with mCRPC, 25 observed the nearest neighbor to one of two prostate cancer cell lines present in the cohort of 452 cancer cell lines (left). The recurrent therapies that nearest neighbors were sensitive to are not widely considered in prostate cancer (right).

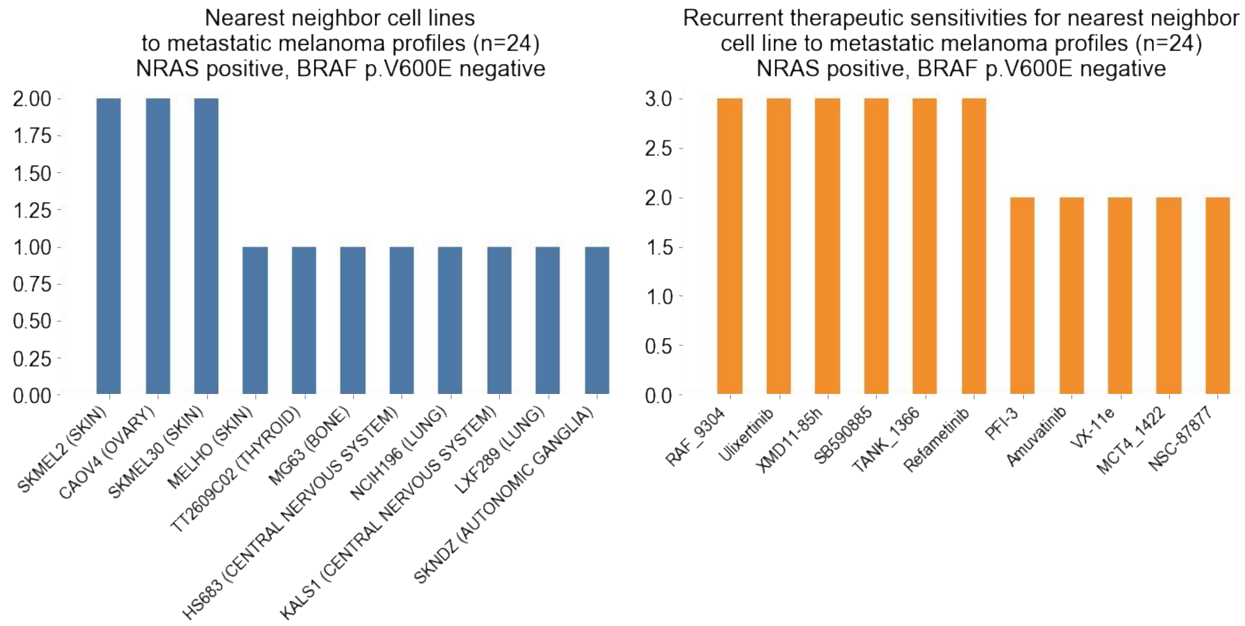


**Supplementary Figure 5.** Patient-to-cell line matchmaking of 24 *NRAS* positive, *BRAF* p.V600E negative metastatic melanoma profiles.

Patient-to-cell line matchmaking was applied to the molecular profiles of 24 patients with metastatic melanoma which harbored *NRAS* genomic alterations but were negative for *BRAF* p.V600E. Recurrent nearest neighbors are observed (left), the top three most common include two skin cancer cell lines and one ovarian cancer cell line for two patient profiles each. Recurrent therapies which nearest neighbors were sensitive to included refametinib (MEK inhibition, 3 patients), ulixertinib (ERK, 3), SB590885 (BRAF inhibitor, 3), and RAF_9304 (RAF, 3).

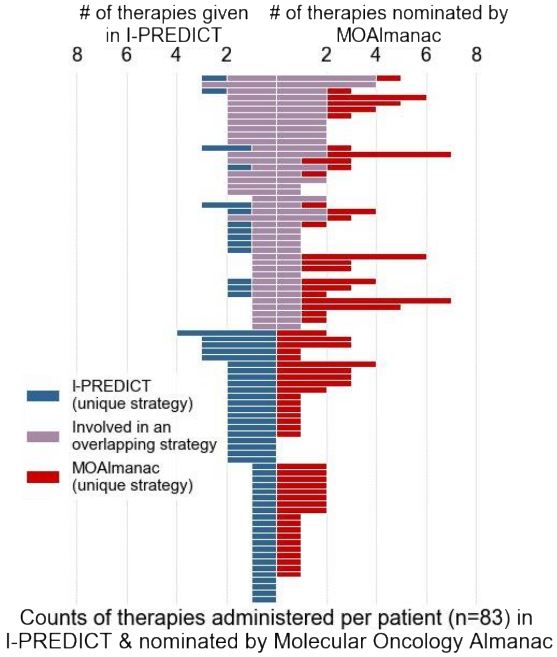


**Supplementary Figure 6.** Comparison between MOAlmanac's nomination and I-PREDICT administered therapies.

For each patient, the number of therapies administered in I-PREDICT are shown (left, blue) and the number of sensitive therapies highlighted by MOAlmanac (right, red). Therapies and associated molecular features targeted were mapped to therapeutic strategies for those administered in I-PREDICT and those highlighted by MOAlmanac (center, purple). A shared therapeutic strategy was observed in 38 (46%) of patients.

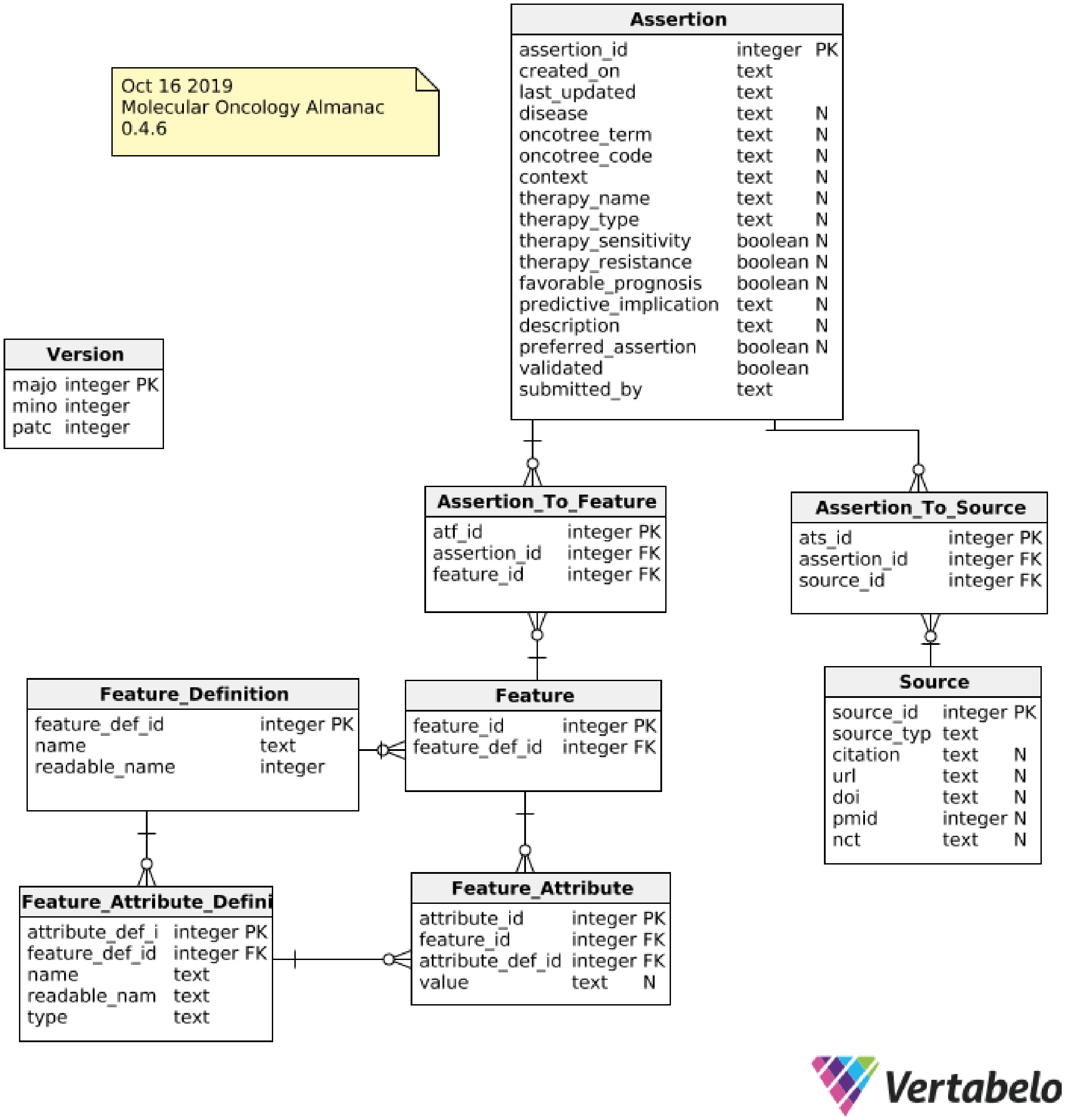


**Supplementary Figure 7.** SQL database schema of the Molecular Oncology Almanac.

The underlying database of MOAlmanac uses SQL and separates records primarily into features, assertions, and sources. Figure generated with Vertabelo.

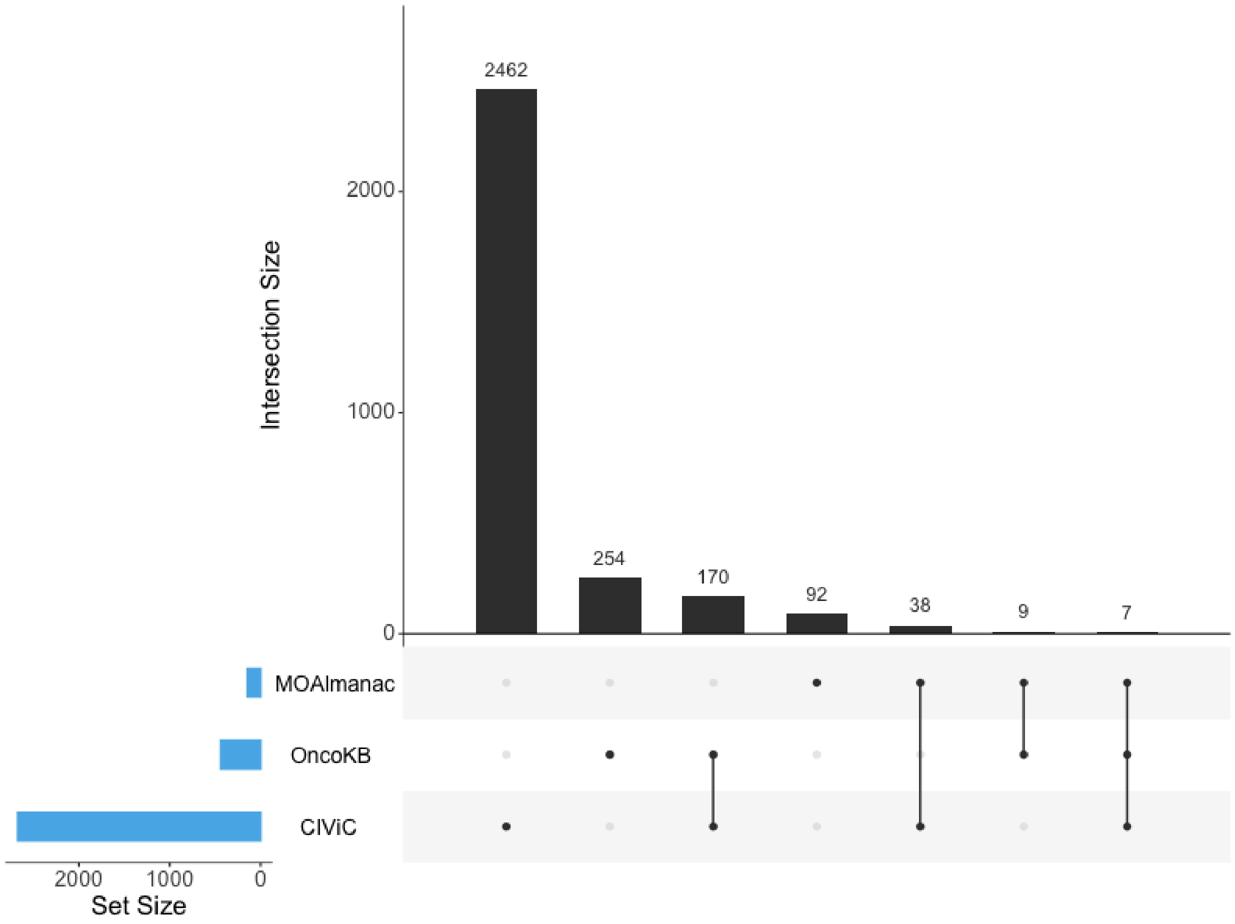


**Supplementary Figure 8.** PubMed ID comparison to OncoKB and CIViC.

UpSet plot comparing PubMed IDs (PMIDs) catalogued by CIViC, Molecular Oncology Almanac, and OncoKB[3,4]. Molecular Oncology Almanac also catalogues clinical guidelines and FDA approvals without PMIDs. No one knowledge base subsumes another. The x-axis represents different intersection sizes, or portions of a venn-diagram. The top row corresponds to PMIDs contained by Molecular Oncology Almanac, the second row OncoKB, and third row CIViC. Colored dots along a knowledge base’s row indicate membership of the intersection displayed along the vertical; for example, the second vertical displays a set of 254 PMIDs which are unique to OncoKB and the third vertical displays a set of 170 PMIDs that are shared by OncoKB and CIViC.

**Supplementary Table 1.** Observed counts of feature types by evidence in retrospective cohorts.

Feature type counts by evidence level for metastatic melanoma (MEL, n = 110) and metastatic castration-resistant prostate cancer (SU2C, n = 150) for relationships suggesting therapeutic sensitivity (sensitive), resistance (resistance), and prognosis (prognosis). Germline event counts have been redacted.

**Supplementary Table 2.** Sensitivity of catalogued relationships in cancer cell lines.

Relationships between genomic alterations and therapeutic sensitivity are evaluated in cancer cell lines with Mann-Whitney-Wilcoxon test (Methods). Of the 44 therapies evaluated, 12 show efficacy for at least one genomic feature.

**Supplementary Table 3.** Comparison between I-PREDICT and MOAlmanac.

Annotated tables of therapies administered by I-PREDICT and highlighted by MOAlmanac (sheet labeled as therapies annotated), molecular features characterized and targeted in I-PREDICT and annotations by MOAlmanac (sheet labeled as features annotated), and sources cited by I-PREDICT categorized by evidence (sheet labeled as citations annotated).

**Supplementary Table 4.** Comparison of features between MOAlmanac, OncoKB, and CIViC.

**Supplementary Table 5.** Comparison of various models that were tested for matchmaking.

Mean average precision, average precision at rank 1-5, and model description for all models used in study to assess genomic similarity between cancer cell lines for matchmaking.

**Supplementary Table 6.** Evaluation of significance of all models.

Pairwise significance testing of all models evaluated. Our best performing model SNF: CGC & FDA was within the noise range of two other models, a multi-pass sort of first using agreement based measure of molecular features associated with an FDA approved therapy followed by agreement based sort of CGC genes mutated by any feature type (Multi-pass sort: FDA & CGC, p=0.3695) and sorting cell lines by their mutant and wild type status of variants in order based on the somatic heuristic in MOAlmanac (Somatic tree, p=0.4765); however, SNF: CGC & FDA observed a stronger AP @ k = 1 in both cases, 0.194 versus 0.167 and 0.119, respectively.
